# Iterative structural homology search identifies new substrates of the protein *O*-fucosyltransferases POFUT3 and POFUT4

**DOI:** 10.64898/2026.05.13.724420

**Authors:** Benjamin Eberand, Huilin Hao, Michelle Cielesh, Kiran Muthukrishnan, Lucas Kambanis, Anthony Ayoub, Yvonne Kong, Jemma Fenwick, Leonie Heilbronn, Richard J. Payne, Freda H Passam, Robert S. Haltiwanger, Mark Larance

## Abstract

*O*-fucosylation plays an essential role in controlling protein folding, secretion and protein-protein interactions within the extracellular space. Recently, we identified a new form of protein *O*-fucosylation occurring on the N-terminal Elastin Microfibril Interaction (EMI) domain of several secreted proteins, mediated by two previously uncharacterized protein *O*-fucosyltransferases, POFUT3 (FUT10) and POFUT4 (FUT11). As all POFUT enzymes (POFUT1-4) are highly specific for the three-dimensional (3D) structure of their substrate protein domains, we postulated that structural homologues of these domains in other proteins may also be *O*-fucosylated. Here, we employed iterative protein structural homology searches as a novel strategy for identifying EMI-like domains that may serve as potential substrates for POFUT3/4. We discovered that microfibrillar-associated protein 2 and 5 (MFAP2/MFAP5) contain EMI-like domains and are *O*-fucosylated at high stoichiometry in human tissues. Unexpectedly, we showed that only POFUT3 is both necessary and sufficient for MFAP2/MFAP5 *O*-fucosylation, despite POFUT4 also having strong protein-protein interactions with MFAP2/MFAP5. Finally, we determined that *O*-fucosylation of MFAP2/MFAP5 is required for their efficient secretion, similar to other EMI domain-containing proteins. Together, these data demonstrate the power of sensitive structural homology analysis in identifying new enzyme-substrate relationships and protein-protein interactions.

## Introduction

Protein *O*-fucosylation is an essential post-translational modification (PTM) that modulates a broad range of physiological and pathological processes such as cellular communication and signal transduction^1,2^. *O*-fucose is present on specific cysteine-rich domains of secreted and cell surface proteins and has key functions in facilitating protein folding, protein-protein interactions and secretion^3–5^. This modification is mediated by protein *O*-fucosyltransferase (POFUT) enzymes, which directly transfer the L-fucose sugar from guanosine diphosphate (GDP)-fucose to protein serine/threonine residues in the endoplasmic reticulum (ER), in a protein domain-specific manner^5–7^. The 3D structure of each POFUT enzymatic pocket is highly conserved across evolution and each shows distinct complementarity to the surface structure of their cognate protein domain substrates. For example, POFUT1 is known only to bind and modify Epidermal Growth Factor-like (EGF-like) repeats containing the motif C^2^XXXX[S/T]C^3^ (where C^2^ and C^3^ represent the second and third cysteine in the EGF), while POFUT2 modifies only Thrombospondin Type 1 Repeats (TSRs) with a C^1−2^XX[S/T]C^2−3^ consensus sequence^8,9^. These substrate domains each contain six conserved cysteines forming three disulfide bonds and are *O*-fucosylated only when these disulfide bonds form correctly. As such, it is hypothesised that POFUT enzymes may play a role in non-canonical protein-folding quality control within the ER, wherein the addition of *O-*fucose may act as a signal indicating proper folding prior to secretion^10,11^. Dysregulation of *O*-fucosylation is associated with severe developmental defects and various cancers, and whole-body knockout of either POFUT1 or POFUT2 in mice is embryonic lethal^12–15^.

Recently, we identified a new form of protein *O*-fucosylation on the N-terminal Elastin Microfibril Interaction (EMI) domain of several extracellular matrix proteins, including multimerin-1 (MMRN1), multimerin-2 (MMRN2) and EMI domain-containing protein 1 (EMID1)^5,16^. We demonstrated that loss of *O*-fucose on the EMI domain reduces protein secretion, highlighting the functional importance of this modification. EMI domain *O*-fucosylation is mediated by two previously uncharacterized protein *O*-fucosyltransferases, POFUT3 (FUT10) and POFUT4 (FUT11)^5^. Like other POFUTs, these enzymes are localised to the ER and only modify correctly folded EMI domains, indicating they also have a role in the ER non-canonical quality control pathway.

Identification of new substrates for the POFUTs has previously been performed through primary protein sequence homology using either BLAST-P analysis, or detection of the substrate consensus motif ^1,17^. However, it is possible that other POFUT substrates exist that only share 3D structural homology and lack strong primary protein sequence homology. Structural homology analysis is a powerful tool for the comparison of protein 3D structures, but until recently has been plagued by slow analysis speed and a lack of comprehensive protein structure databases to search^18^. However, the complementary development of the Foldseek rapid structural homology analysis algorithm and databases containing AlphaFold structural predictions of all human proteins has enabled rapid proteome-wide structural searches^19^.

In this study, we aimed to explore the diversity of POFUT3/4 substrates. As POFUTs are highly specific for certain substrate structures, we hypothesised that any proteins containing domains structurally homologous to an EMI domain may also be *O*-fucosylated. Here, we employed an iterative Foldseek search to identify EMI-like protein domains, which may represent novel targets for POFUT3/4-mediated *O*-fucosylation. We show that the structure of the matrix binding domain (MBD) in the microfibrillar-associated protein 2 and 5 (MFAP2/MFAP5) is highly homologous to the EMI domain in 3D structure but shares low homology in primary sequence. These two extracellular matrix accessory proteins share high sequence similarity and are known to regulate TGF-β signalling associated with fibrillin microfibrils^20^. We demonstrate that both MFAP2 and MFAP5 are *O*-fucosylated within their MBDs in mammalian cells and tissues. Using co-immunoprecipitation assays and knockout cell lines, we show that MFAP2 and MFAP5 bind with high affinity to both POFUT3 and POFUT4. However, only POFUT3 is necessary and sufficient for their *O*-fucosylation. Furthermore, blocking MBD *O*-fucosylation significantly reduced the secretion of MFAP2 and MFAP5, suggesting a conserved role for *O*-fucose in protein-folding quality control. Together, these findings demonstrate that structural homology analysis can accurately identify novel substrates of domain-specific enzymes and enables expansion of the substrate repertoire of POFUT3/4.

## Results

### Iterative Foldseek search strategy identifies new *O*-fucosylation substrates for POFUT3/4

To identify new substrate domains for *O*-fucosylation by POFUT3/4, we generated a predicted structure of the EMI domain of human multimerin-1 (MMRN1) in Colabfold^21^, which is a well-established POFUT3/4 substrate^5,16^. We used Foldseek to align this structure against the entire human AlphaFold Protein Structure Database (AFDB50), identifying structural homologues that consisted largely of other *bona fide* EMI domain-containing proteins (**Fig. 1a**, grey boxes). The EMI-like domains from this first iteration were then used individually for additional Foldseek searches in an iterative process to generate a protein homology network (**Fig. 1a**, and **Supplementary Table 1**). From this, several proteins were identified as containing highly structurally homologous EMI-like domains, despite lacking a *bona fide* EMI domain (**Fig. 1a**, red-boxed proteins). This included the matrix binding domain (MBD) of microfibrillar-associated proteins 2 and 5 (MFAP2/MFAP5) (**Fig. 1b**), and several other extracellular matrix proteins such as SNED1 (sushi, nidogen and EGF-like domain-containing protein 1) and VWA2 (von Willebrand factor A domain-containing protein 2). Additional matches with lower homology were observed, however manual inspection of their structures showed poor structural alignment (**Fig. 1a**, grey dashed, white box proteins). The multiple protein structure alignment tool FoldMason^22^ was used to generate a primary sequence alignment for all EMI and EMI-like domains that were identified (**Fig. 1c**). This showed limited primary sequence homology between EMI and EMI-like domains with only partial alignment of key cysteine residues, particularly lacking the characteristic “CC” in the C-terminal half of the EMI domain^23^. However, the threonine and serine residues in the N-terminal half of all EMI-like domains, which are known *O*-fucosylation sites in EMI domains, were conserved (**Fig. 1c**, red triangle). In addition, several residues in the C-terminal half of the domains were also highly conserved across EMI and EMI-like domains and may play important roles in protein-protein interactions (**Fig. 1c**, asterisks).

**Figure 1.**
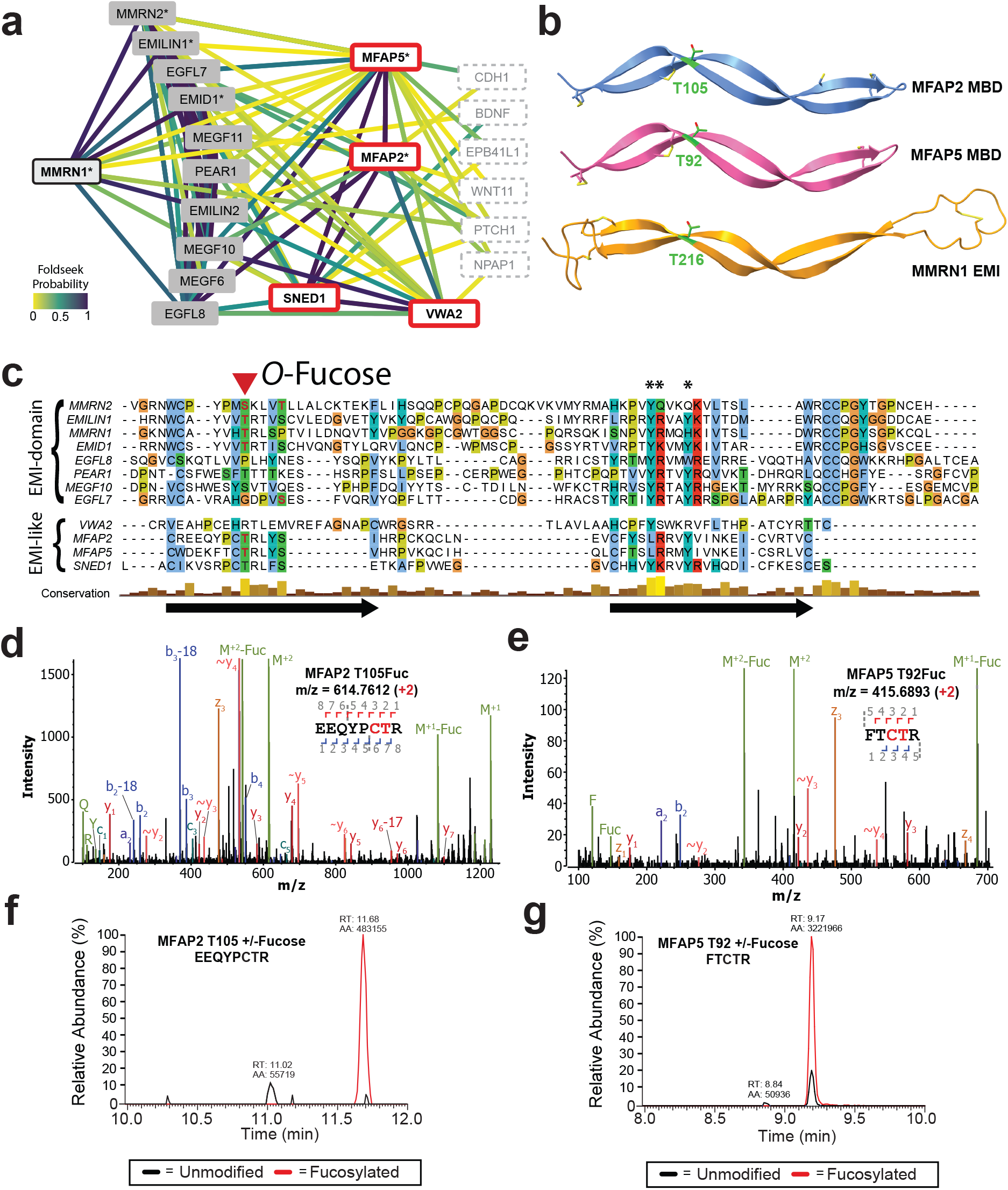
Iterative Foldseek search strategy identifies new *O*-fucosylation substrates in mammals. **a**, Structural homology network generated aligning the EMI domain of human MMRN1 (residues 207-282) against the human AFDB50 database using an iterative Foldseek search. Lines connecting proteins are coloured according to their predicted structural homology confidence. Proteins known to contain an EMI domain are represented in grey boxes, while proteins without an annotated EMI domain are indicated by red boxes. Proteins in the dashed boxes represent dubious results. Asterisks are *O*-fucosylated proteins. **b**, AlphaFold2 structures for the matrix binding domain (MBD) of human MFAP2 (residues 93-143) and MFAP5 (residues 84-130), structurally aligned with the EMI domain (residues 207-282) of human MMRN1. The conserved site of *O*-fucosylation is highlighted in green, and key disulfide bonds are shown in yellow. **c**, FoldMason sequence alignment of all known EMI domain and EMI-like domain containing proteins, coloured by residue type. The conserved putative *O*-fucosylation site is demonstrated by a red triangle, with experimentally confirmed *O*-fucosylation sites highlighted in red text. Other key conserved residues are indicated by asterisks. **d**, Annotated EAD MS/MS spectra for the *O*-fucosylated human MFAP2 peptide, derived from human adipose tissue. Human adipose tissue was digested with trypsin and analysed with targeted LC-MS/MS using EAD fragmentation. Ions are annotated as a-ions, b-ions, c-ions, y-ions, z-ions, ~y-ions with a neutral loss (such as a loss of fucose), M (intact precursor), M-Fuc (intact precursor with neutral loss of fucose) or Fuc (fucose oxonium ion). **e**, Annotated EAD MS/MS spectra for the *O*-fucosylated human MFAP5 peptide, derived from human adipose tissue. **f**, Extracted ion chromatograms (EICs) of different glycoforms of the peptide containing the MFAP2 T105 *O*-fucose site in human adipose tissue. **g**, EICs of different glycoforms of the peptide containing the MFAP5 T92 *O*-fucose site in human adipose tissue.

The MBD of human MFAP2/MFAP5 displayed strong structural homology to MMRN1 EMI domain, with root mean square deviation (RMSD) values of 0.82 Å and 1.17 Å across aligned regions from matchmaker analysis in ChimeraX, respectively (**Fig. 1b**). RMSD values less than 2 Å are considered high homology^24^. Crucially, multiple structure alignment indicated that the MBDs of MFAP2 and MFAP5 each possess a putative *O*-fucosylation site at a position and orientation equivalent to that of the MMRN1 EMI domain, specifically residue T105 in MFAP2 and T92 in MFAP5 (**Fig. 1b**, green residues). Hence, we hypothesised these sites may be *O*-fucosylated by POFUT3/4. Analysis of the primary sequence surrounding these sites in MFAP2 and MFAP5 showed 100% conservation across vertebrate evolution (**Supplementary Fig. 1**), suggesting these sites are important for protein function. To determine if MFAP2 and MFAP5 are endogenously *O*-fucosylated, human white adipose tissue, where these proteins are highly expressed (**Supplementary Fig. 2**), was analysed by targeted glycoproteomic analysis. This analysis confirmed *O*-fucosylation of MFAP2 at T105 and MFAP5 at T92 (**Fig. 1d,e**). Further, extracted ion chromatograms (EICs) for both the *O*-fucosylated and unmodified glycoforms of these sites revealed this modification to occur at high stoichiometry (>90%) in human white adipose tissue (**Fig. 1f,g**). It should be noted that these small *O*-fucosylated peptides like that of MFAP5 undergo significant neutral loss of fucose through in-source decay.

**Figure 2.**
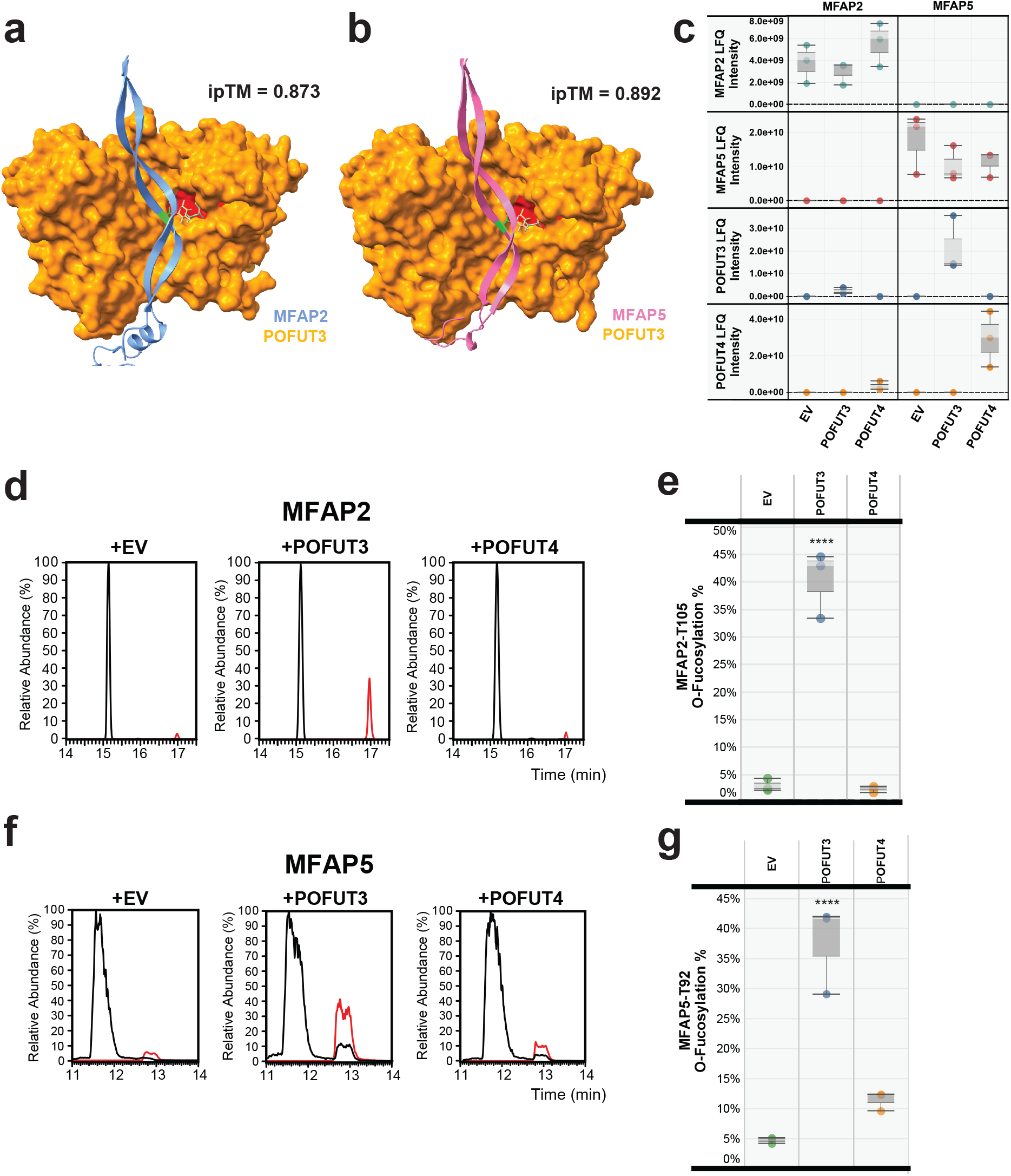
MFAP2 and MFAP5 bind both POFUT3 and POFUT4 at high affinity but prefer *O*-fucosylation via POFUT3. **a**, AlphaFold2-multimer predicted structure for the interaction of human POFUT3 (residues 81-479, orange), with the matrix binding domain (MBD) and C-terminal portion of human MFAP2 (residues 97-183, blue). The site of *O*-fucosylation is highlighted in green, while the active site of the enzyme is highlighted in red. GDP-fucose is shown in yellow stick. The interface predicted template modelling (ipTM) score indicates confidence in the predicted structure of this interaction, where a score ≥0.8 is considered very high confidence. **b**, AlphaFold2-multimer predicted structure for the interaction of human POFUT3 (residues 81-479, orange), with the MBD and C-terminal portion of human MFAP5 (residues 84-173, pink). **c**, Boxplots showing label-free quantification (LFQ) intensity of enzyme and substrate proteins immunoprecipitated with either MFAP2-FLAG, or MFAP5-FLAG. Proteins were isolated from lysates of HEK293 cells transiently transfected with MFAP2-FLAG, MFAP5-FLAG or an empty vector (EV), and co-transfected with either POFUT3, POFUT4, or an empty vector (EV) (n = 3). **d**, EICs of different glycoforms of the peptide containing the MFAP2 T105 *O*-fucose site in HEK293 cell lysates, analysed by LC-MS/MS. Red lines, *O*-fucose modified; black lines, unmodified. **e**, Boxplot showing the relative abundance of MFAP2 substrate peptide that is *O*-fucosylated in HEK293 cell lysates. **f**, EICs of different glycoforms of the peptide containing the MFAP5 T92 *O*-fucose site in HEK293 cell lysates. Red lines, *O*-fucose modified; black lines, unmodified **g**, Boxplot showing the relative abundance of MFAP5 substrate peptide that is *O*-fucosylated in HEK293 cell lysates. Statistical analysis was performed with unpaired, two-tailed t-test in Prism 7. ****P < 0.0001. All data are shown as mean ± s.d. from biological triplicates of three individual transfections (**Supplementary Fig. 6**).

### MFAP2/MFAP5 bind both POFUT3 and POFUT4 but prefer *O*-fucosylation by POFUT3

The previously known EMI domain substrates of POFUT3/4 were shown to have strong protein-protein interactions with both enzymes^5^. To examine if MFAP2 and MFAP5’s MBDs also bind POFUT3/4, we analysed the interactions between these putative substrates and POFUT3/4 using AlphaFold2-multimer in Colabfold. In this analysis the MBD of either MFAP2, or MFAP5, was folded alongside each human fucosyltransferase (FUT1-9 and POFUT1-4). High confidence interactions with an interface predicted template modeling (ipTM) score >0.8 were identified only when the MBD was folded with POFUT3 (**Fig. 2a,b**) and POFUT4 (**Supplementary Fig. 3-5**). POFUT3 showed ipTM scores of 0.873 and 0.892 for MFAP2 and MFAP5 respectively, while POFUT4 showed slightly lower ipTM scores of 0.782 and 0.829. This suggests that the MBD may bind more strongly to POFUT3 than POFUT4, a preference not observed for any EMI-domain to date^5^. Importantly, in the AlphaFold2-multimer predicted structures of MFAP2 and MFAP5’s MBDs bound to POFUT3/4, T92 and T105 were the residues closest to the enzyme active site and thus may be *O*-fucosylated. To validate this predicted interaction, co-immunoprecipitation mass spectrometry analysis was performed^5^. HEK293T cells were transiently transfected with mammalian expression plasmids encoding FLAG-tagged MFAP2 or MFAP5, in either the presence or absence of GFP-tagged POFUT3, or POFUT4. Anti-FLAG co-immunoprecipitation of MFAP2/MFAP5 showed significant co-isolation of POFUT3 and POFUT4 for both substrates in cell lysates, consistent with the binding predicted by AlphaFold2-multimer (**Fig. 2c**). Additionally, targeted glycoproteomic analysis was used to estimate the stoichiometry of *O*-fucosylation in these co-immunoprecipitations. This showed that *O*-fucosylation of both MFAP2 and MFAP5 was significantly increased when co-transfected with POFUT3 compared with empty vector. However, co-transfection with POFUT4 showed either very little or no significant *O*-fucosylation of either protein (**Fig. 2d-g, Supplementary Fig 6**), suggesting a preference for the EMI-like MBDs of MFAP2/MFAP5 to be *O*-fucosylated by POFUT3.

**Figure 3.**
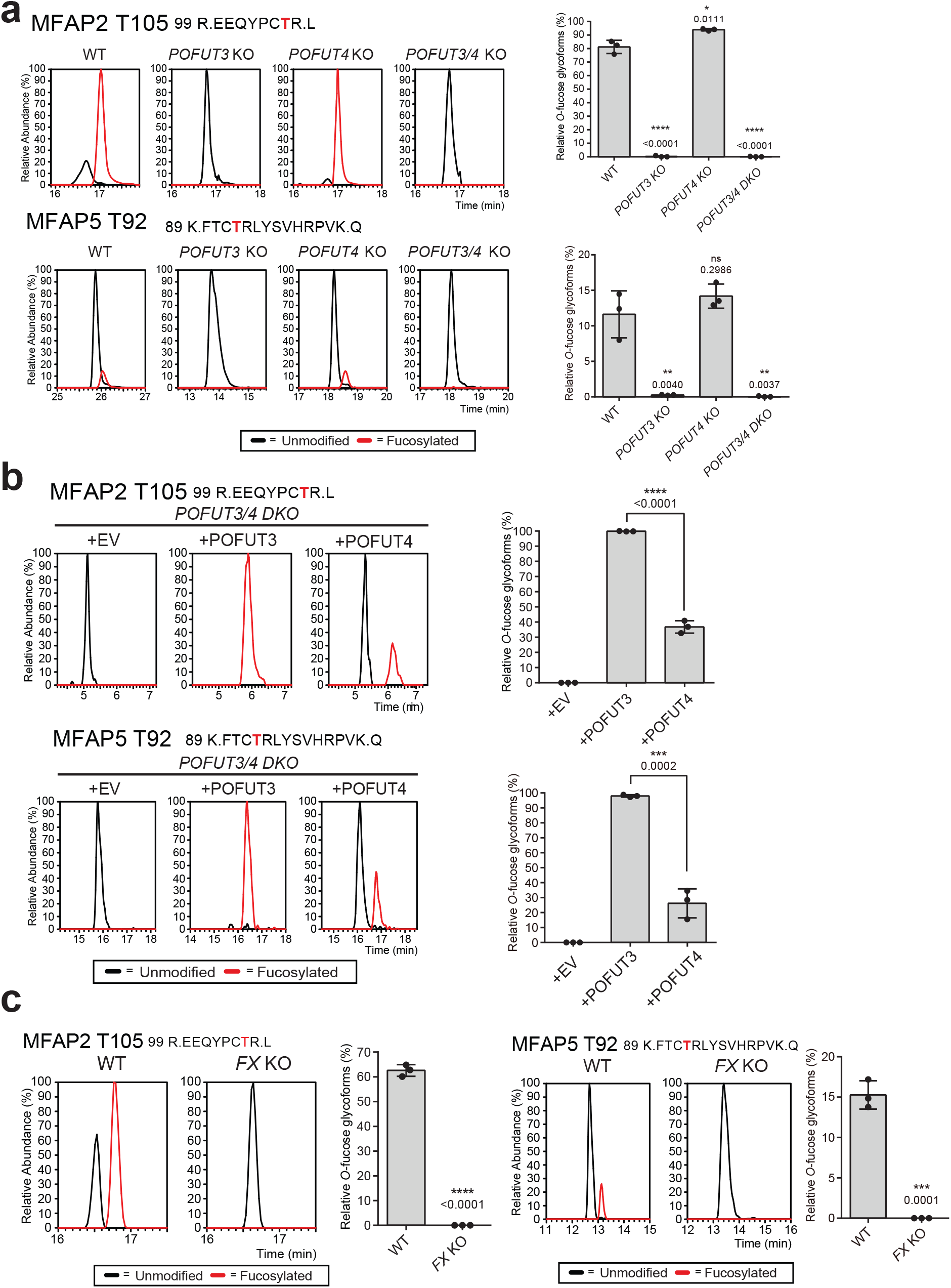
POFUT3 alone is sufficient for *O*-fucosylation of both MFAP2 and MFAP5. **a**, EICs of different glycoforms of MFAP2/MFAP5 *O*-fucose site-containing peptides, produced in HEK293T wild-type (WT), *POFUT3* KO, *POFUT4* KO, or *POFUT3/4* DKO cells. Red lines represent the relative abundance of *O*-fucosylated peptide while the black lines represent the abundance of unmodified peptide. Statistical analysis was performed with unpaired, two-tailed t-test in Prism 7. *P < 0.1; **P < 0.01; ****P < 0.0001 compared to control (WT cells). **b**, EICs of the different glycoforms of the MFAP2/MFAP5 *O*-fucose site-containing peptides produced in HEK293T *POFUT3/4* DKO cells that co-transfected with plasmids encoding POFUT3, POFUT4, or an empty vector (EV). Red lines represent the relative abundance of *O*-fucosylated peptide while the black lines represent the abundance of unmodified peptide. Statistical analysis was performed with unpaired, two-tailed t-test in Prism 7. ***P < 0.001; ****P < 0.0001. **c**, EICs of different glycoforms of MFAP2/MFAP5 *O*-fucosylated peptide produced in either HEK293T WT or *FX* KO (lacking GDP-fucose synthesis) cells. Red lines represent the relative abundance of *O*-fucosylated peptide while the black lines represent the abundance of unmodified peptide. Statistical analysis was performed with unpaired, two-tailed t-test in Prism 7. ***P < 0.001; ****P < 0.0001 compared to control (WT cells). All data are shown as mean ± s.d. from biological triplicates of three individual transfections (**Supplementary Fig. 7**).

### POFUT3 is necessary for *O*-fucosylation of both MFAP2 and MFAP5 in cells

To determine if POFUT3 is the main enzyme responsible for *O*-fucosylation of MFAP2 and MFAP5, we employed *POFUT3* and *POFUT4* single and double knockout (KO) HEK293T cell lines^5^. Each cell line was transfected with MFAP2/MFAP5 6xHis-tagged expression plasmids and the conditioned culture media was harvested after three days of incubation. The secreted 6xHis-tagged MFAP2/MFAP5 proteins were purified and analysed with glycoproteomic site-mapping. Both MFAP2-T105 and MFAP5-T92 were observed to be *O*-fucosylated in wildtype cell culture media and completely lost the *O*-fucose in *POFUT3/4* double KO cell culture media (**Fig. 3a, Supplementary Fig. 7a**), consistent with our previous observations for EMI-containing proteins^5^. Interestingly, complete loss of *O*-fucose was also observed in *POFUT3* single KO cells, while knocking out *POFUT4* had no significant impact on the *O*-fucosylation levels for MFAP2/MFAP5. Such substrate preference between POFUT3 and POFUT4 was not observed in any of the previously tested EMI-containing proteins^5^. Moreover, transient transfection of a POFUT3 expression plasmid in *POFUT3/4* double KO cells fully restored the *O*-fucosylation in both MFAP2/MFAP5 and drove the stoichiometry to near 100% (**Fig. 3b, Supplementary Fig. 7a**). This suggests that POFUT3 alone is necessary for MFAP2/MFAP5 *O*-fucosylation. Interestingly, transfection of a POFUT4 expression plasmid in *POFUT3/4* double KO cells partially rescued the *O*-fucosylation of MFAP2/MFAP5 with ~30% stoichiometry (**Fig. 3b, Supplementary Fig. 7b**), indicating POFUT4 may have some ability to *O*-fucosylate the MBD when added in excess by over-expression. We confirmed that GDP-fucose synthesis is required for the *O*-fucosylation of MFAP2/MFAP5 by testing the modification status of these EMI-like substrates expressed in HEK293T cells with a null mutation in GDP-L-fucose synthase (GFUS/FX)^5^. This showed complete loss of *O*-fucose for MFAP2/MFAP5 in *FX* KO cells (**Fig. 3c, Supplementary Fig. 7c**).

### POFUT3 alone is sufficient for *O*-fucosylation of MFAP2 MBD in vitro

To determine whether POFUT3 can efficiently *O*-fucosylate the MFAP2 MBD, we performed enzymatic assays on purified, non-fucosylated MFAP2 using recombinant GFP-POFUT3/4. GFP-POFUT3/4 were purified from HEK293F cells using POFUT3/4 expression constructs^25^, while non-fucosylated MFAP2 was purified from HEK293T *FX* KO cells adapted to serum-free suspension culture^26^. MFAP2 was incubated with either GFP-POFUT3 or GFP-POFUT4, alongside GDP-fucose for up to 4 hours at 37°C, then analysed by glycoproteomics. The same incubation and analysis were also performed upon purified MMRN1 N-terminal EMI to compare modification efficiency with a known substrate^5^. While POFUT3 was able to add fucose to MFAP2-T105 in a time-dependent manner and approached saturation around 4 hours, no modification by POFUT4 was detected over the same time course (**Fig. 4a**,**b**). In contrast, N-terminal EMI domain showed no such enzyme preference at either of its modification sites (T216, T265) (**Supplementary Fig. 8a**,**b**). This enzymatic assay was also repeated with MFAP2 produced from SHuffle *E. coli* (**Supplementary Fig. 9a-c**), a bacterial strain specialised for production of disulfide bonded proteins^27^, which showed similar results albeit with lower overall modification stoichiometry (**Supplementary Fig. 9d**). These results indicate that POFUT3 is required for *O*-fucosylation of MFAP2-T105.

**Figure 4.**
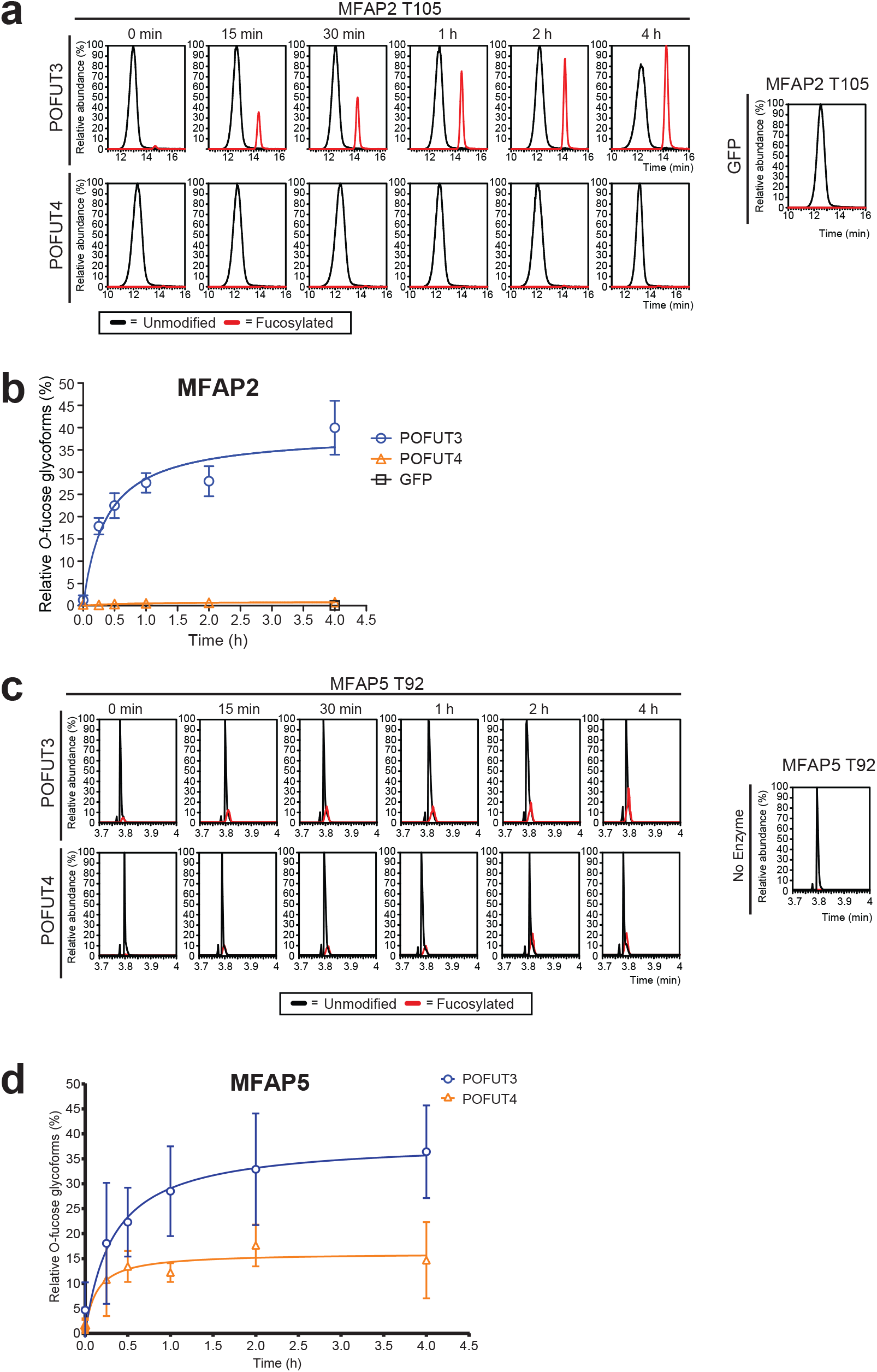
MFAP2 shows a preference for *O*-fucosylation by POFUT3. **a**, 0.1 μM purified GFP-POFUT3, GFP-POFUT4 or GFP (negative control) was incubated with 0.5 μM non-fucosylated MFAP2, 100 μM GDP-fucose and 300 μM MnCl2 at 37°C for up to four hours. Reaction products were reduced, alkylated, digested with trypsin and analysed by nano LC-MS/MS. EICs of peptides containing the T105 *O*-fucose site were examined at the indicated time points to compare relative abundance of the modified and unmodified peptide. A significant preference for POFUT3 is observed, as POFUT4 shows no modification after four hours. Red lines represent the relative abundance of the *O*-fucosylated peptide while the black lines represent the abundance of the unmodified peptide. **b**, Relative abundances of *O*-fucosylation for each reaction were calculated from the EICs of peptides containing the T105 *O*-fucose site and plotted as time-dependent curves. Data are presented as mean ± s.d. from biological triplicates. **c**, 0.5 μM non-fucosylated synthetic MFAP5 MBD was incubated with 0.1 μM purified GFP-POFUT3, GFP-POFUT4 or no enzyme (negative control) alongside 100 μM GDP-fucose and 300 μM MnCl2 at 37oC for up to four hours. Reaction products were reduced, alkylated, digested with trypsin and analysed by nano LC-MS/MS. EICs of peptides containing the T92 *O*-fucose site were examined at the indicated time points to compare relative abundance of the modified and unmodified peptide. Red lines represent the relative abundance of the *O*-fucosylated peptide while the black lines represent the abundance of the unmodified peptide. **d**, Relative abundances of *O*-fucosylation for each reaction were calculated from the EICs of peptides containing the T92 *O*-fucose site and plotted as time-dependent curves. Data are presented as mean ± s.d. from biological triplicates.

We also performed this enzymatic assay for the non-fucosylated MFAP5 MBD produced synthetically by native chemical ligation of synthesised peptides, followed by a folding reaction to generate disulfide bonds and subsequent purification (**Supplementary Fig. 10, 11**). This synthetic MFAP5 was incubated with either GFP-POFUT3, GFP-POFUT4 or no enzyme, alongside GDP-fucose for 4 hours at 37°C, then analysed by glycoproteomics. POFUT3 was able to add fucose to MFAP5-T92 in a time dependent matter and reached saturation at roughly 35% modification after 4 hours (**Fig. 4c,d**). Interestingly, POFUT4 was also able to add fucose to MFAP5-T92, albeit with reduced efficiency, reaching saturation at roughly 15% modification. These results indicate that while POFUT3 is the preferred enzyme for both MFAP2-T105 and MFAP5-T92 *O*-fucosylation, POFUT4 is also able to *O*-fucosylate MFAP5 when added in excess.

### *O*-fucosylation is necessary for the secretion of MFAP2 and MFAP5

As all human POFUTs are known to exhibit regulatory effects upon protein secretion^5,16,28,29^, we wanted to determine if *O*-fucosylation had a similar effect on MFAP2 and MFAP5. We performed secretion assays for FLAG-tagged MFAP2/MFAP5 in HEK293T cells, as described previously^5^. Expression plasmids encoding alanine mutants at either T105 for MFAP2, or T92 for MFAP5 were used alongside wildtype constructs for analysis of secretion effects, relative to an IgG secretion positive/loading control. We found that when *O*-fucosylation of MFAP2 was blocked by mutation of the site to alanine, secretion was significantly reduced by >50% (**Fig. 5a,b**). However, analysis of the T92A MFAP5 mutant showed no significant impact upon secretion (**Fig. 5c,d**). To confirm this result, we performed MFAP2/MFAP5 secretion assays in *POFUT3/4* double KO and *FX* KO HEK293T cell lines. Expression plasmids encoding wildtype MycHis-tagged MFAP2, MFAP5, or MMRN1 N-terminal EMI were transfected into *POFUT3/4* double KO, *FX* KO and wildtype (WT) HEK293T cell lines, alongside the IgG secretion control. *POFUT3/4* double KO led to a 50-80% reduction in both MFAP2 and N-terminal EMI secretion (**Fig. 5e,g**). A similar reduction was observed in the *FX* KO cells, which cannot synthesise GDP-fucose (**Fig. 5f,h**). The effect of *POFUT3/4* double KO on MFAP5 was less severe, with only a 30% reduction in secretion (**Fig. 5i,k**). However, *FX* KO led to an approximate 60% loss of secretion for MFAP5, in line with the reduction observed for N-terminal EMI (**Fig. 5j,l**), indicating that other fucosylation events may also contribute to MFAP5 secretion. Together, these findings suggest that *O*-fucosylation of the EMI-like domains in MFAP2/MFAP5 regulates their secretion, and thus has functional similarity to *O*-fucosylation of the canonical EMI domains.

**Figure 5.**
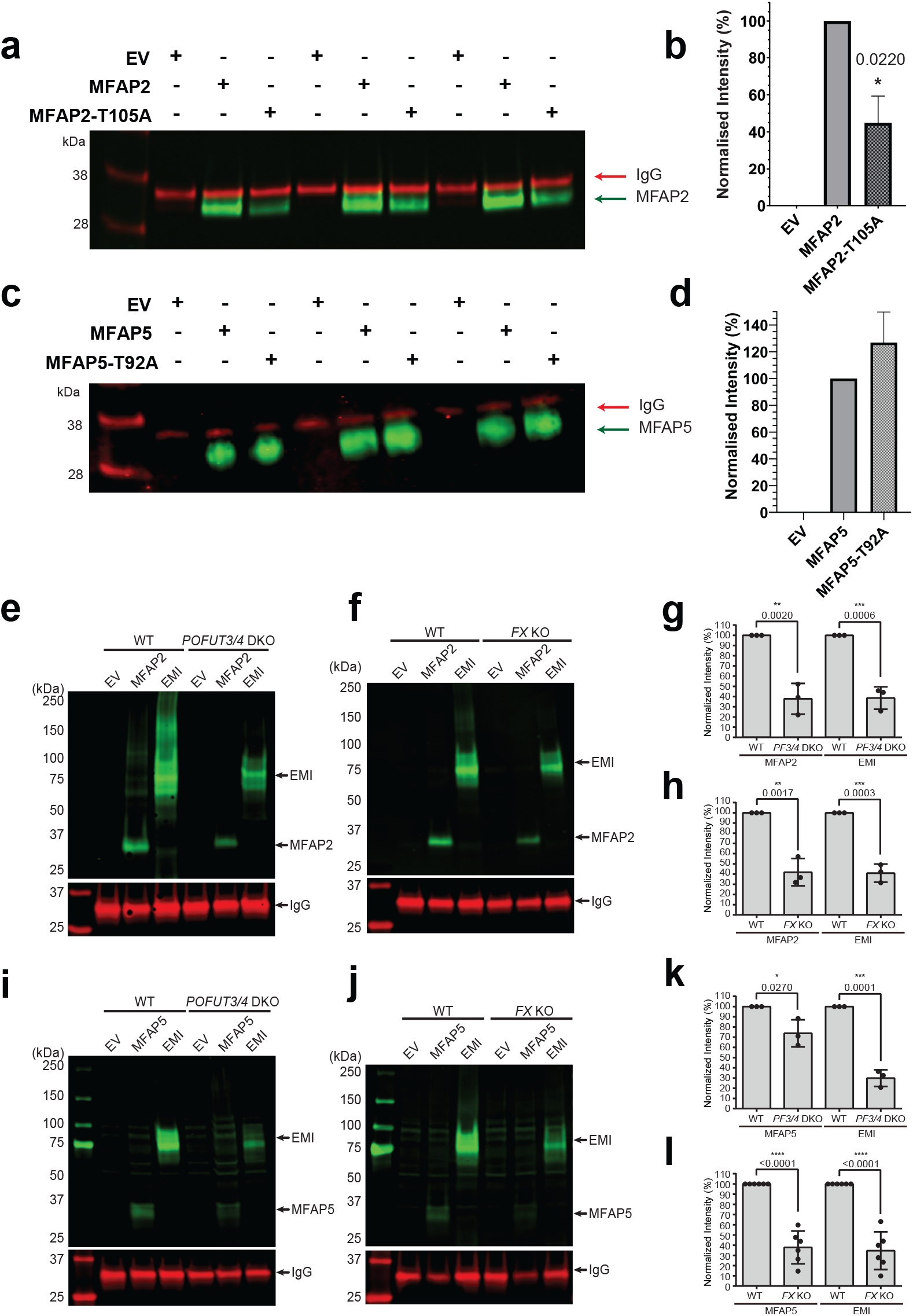
*O*-fucosylation is necessary for the secretion of MFAP2 and MFAP5. **a**, HEK293T WT cells were transfected with plasmids encoding FLAG-tagged MFAP2 WT, MFAP2-T105A, or empty vector (EV), alongside an IgG secretion control. Cells were cultured for 2 days. Culture media was analysed by western blot probed with anti-FLAG and anti-human IgG antibodies. **b**, Bar graphs show normalised band intensity of MFAP2, MFAP2-T105A, and EV. Band intensity was normalised to IgG band intensity. Data is shown as mean ± s.d. from biological triplicates of three individual transfections. Statistical analysis was performed with unpaired, two-tailed t-test in Prism 7. *P < 0.05 compared to MFAP2 WT. **c**, HEK293T WT cells were transfected with plasmids encoding FLAG-tagged MFAP5 WT, MFAP5-T92A, or EV, alongside an IgG secretion control. Cells were cultured for 2 d. Culture media was analysed by western blot probed with anti-MFAP5 and anti-human IgG antibodies. **d**, Bar graphs show normalised band intensity of MFAP5, MFAP5-T92A and EV. Band intensity was normalised to IgG band intensity. Data is shown as mean ± s.d. from biological triplicates of three individual transfections. **e**, HEK293T WT or *POFUT3/4* KO cells were transfected with plasmids encoding Myc-tagged MFAP2, N-terminal EMI (positive control), or EV, alongside an IgG secretion control. Two-day culture media was analysed by western blot probed with anti-Myc and anti-human IgG antibodies. **f**, HEK293T WT or *FX* KO cells were transfected with plasmids encoding Myc-tagged MFAP2, N-terminal EMI, or EV, alongside an IgG secretion control. Two-day culture media was analysed by western blot probed with anti-Myc and anti-human IgG antibodies. **i**, HEK293T WT or *POFUT3/4* KO cells were transfected with plasmids encoding Myc-tagged MFAP5, N-terminal EMI, or EV, alongside an IgG secretion control. Two-day culture media was analysed by western blot probed with anti-Myc and anti-human IgG antibodies. **j**, HEK293T WT or *FX* KO cells were transfected with plasmids encoding Myc-tagged MFAP5, N-terminal EMI, or EV, alongside an IgG secretion control. Two-day culture media was analysed by western blot probed with anti-Myc and anti-human IgG antibodies. **g**,**h**,**k**,**l**, Bar graphs show quantified normalised band intensity obtained in e (g), f (h), i (k) and j (l), respectively. Data are presented as mean ± s.d. from biological triplicates (**Supplementary Fig. 14 and 15**). Statistical analysis was performed with unpaired, two-tailed t-test in Prism 7. *P < 0.1; **P < 0.01; ***P < 0.001; ****P < 0.0001 compared to control (WT cells).

## Discussion

*O*-fucosylation plays an essential role in controlling protein folding, secretion and protein-protein interactions within the extracellular space. Our recent discovery of POFUT3/4 as protein *O*-fucosyltransferases that modify EMI domains highlights the significant gaps in our understanding of the protein *O*-fucosylation landscape. Here, we used iterative protein structural similarity analysis to identify MFAP2 and MFAP5 as new *O*-fucosylated proteins modified with high stoichiometry in human tissue. Unexpectedly, we found a strong preference for POFUT3-mediated modification of MFAP2/MFAP5, which has not previously been observed for known EMI-domain substrates. Our data shows that POFUT3 is necessary and sufficient for MFAP2/MFAP5 *O*-fucosylation, and that *O*-fucosylation of MFAP2/MFAP5 is required for their efficient secretion, similar to other EMI-domain containing substrates. Together, these data demonstrate the power of sensitive structural homology analysis in identifying new enzyme-substrate relationships and protein-protein interactions.

The landscape of *O*-fucosylation has been expanded with the discovery of POFUT3/4-mediated EMI domain *O*-fucosylation^5,16^. The results we report here extend the substrate pool of *O*-fucosylation beyond EMI domains to include the matrix binding domain (MBD) of MFAP2 and MFAP5, which we designate as an “EMI-like” domain. In contrast to all previously examined EMI domains, we observed that the MBD of MFAP2/MFAP5 has a strong preference for *O*-fucosylation by POFUT3. In contrast, POFUT4 only *O*-fucosylated MFAP2/MFAP5 when POFUT4 was over-expressed in the DKO cells. One explanation for this POFUT3-preference may be due to subtle differences in the structure of the enzyme-substrate interface. One notable distinction is within the extended C-terminal region unique to POFUT3 and POFUT4, which we previously hypothesised may contribute to substrate specificity^5^. POFUT4’s C-terminal structure is predicted to be more closed and sterically hindered than that of POFUT3 (**Supplementary Fig. 12a**,**b**) due to the presence of several additional charged residues (E442, E462, K447, R489) likely forming salt bridges (**Supplementary Fig. 12c**,**d**). These salt bridges are absent in POFUT3’s equivalent region, allowing greater flexibility^30^. As a result, the disulfide-linked inflexible structure of MFAP2/MFAP5’s MBD may have destabilising steric clashes with the C-terminal region of POFUT4 (**Supplementary Fig. 12f**,**g**), leading to the observed preference for POFUT3 by these new EMI-like domains. Over a longer course of time or when POFUT4 is in excess, this unfavourable interaction may still occur infrequently, explaining how transfection of abundant POFUT4 was able to partially rescue MFAP2/MFAP5 *O*-fucosylation (**Fig. 3b**). Furthermore, immediately following MFAP2’s MBD is a disulfide-rich ShKt domain, further imposing inflexibility upon the structure, which is absent in MFAP5. This key difference may contribute to the more pronounced preference for POFUT3 observed for MFAP2 in the results of both our co-immunoprecipitations and enzymatic assays (**Fig 2d-g, Fig. 4**).

In contrast to the EMI-like domain structure, true EMI domains feature an extended beta turn structure (**Supplementary Fig. 12e**), imparting flexibility upon the domain and likely allowing less restricted binding to both enzymes. The biological reasons for this preference may be linked to the expression patterns of POFUT3/4 and MFAP2/MFAP5 in different tissues. For example, in the human lung single-cell RNA sequencing database from GTEx^31^, filtering for cells with the MFAP2 mRNA showed ~16% of these cells also contained POFUT3 mRNA (**Supplementary Fig. 13a**), while none of the MFAP2-expressing cells contained POFUT4 mRNA. Similarly, filtering for MFAP5 mRNA shows significantly greater co-localisation with POFUT3 mRNA than POFUT4 mRNA (**Supplementary Fig. 13b**). In general, POFUT3 is highly expressed in stem cells and endothelial cells similar to MFAP2/MFAP5^32^, while POFUT4 is strongly expressed in most other human cell types. Future investigation into *Pofut3 or Pofut4* knockout animal models will be necessary to identify the distinct functions of these enzymes across mammalian tissues and cell types. Furthermore, single cell proteomic analysis of endothelial tissue would help to confirm the observed co-localisation of POFUT3 with MFAP2/MFAP5 at the protein level, addressing the shortfalls of transcriptomics-based co-expression analysis^33^.

Here, we have also shown that secretion of MFAP2/MFAP5 is regulated by *O*-fucosylation, similar to known substrates of other POFUT enzymes. MFAP2 and MFAP5 function in the extracellular matrix as components of fibrillin microfibrils, and are known to directly interact with fibrillin proteins (FBN1 and FBN2) through their matrix binding domain (MBD)^34,35^. Once bound to microfibrils, MFAP2/MFAP5 interact with a wide range of proteins such as von Willebrand factor (vWF)^36^ and transforming growth factor beta (TGF-β)^37^ to mediate their core functions, which are commonly associated with vascular phenotypes from loss of function studies. For example, *Mfap2* knockout in zebrafish exhibited vessel dilation and vascular wall defects^38^, while *Mfap2/Mfap5* double knockout mice studies have demonstrated age-dependent aortic dilation^20^. *Mfap2* knockout mice also showed large vessel haemostasis defects (increased bleeding time) and dysregulation of platelet function^35,39^. In humans, nonsense mutation of human MFAP5 close to the C-terminus (R158*) causes protein truncation and reduced secretion, leading to an increased risk of thoracic aortic aneurysm and aortic dissection disorder^40^, indicating the importance of this protein’s correctly folded structure. This phenotype also has significant overlap with disrupted TGF-β signalling, which perturbs healthy angiogenesis and vascular wall integrity^41,42^. In extracellular samples from human patients^40^, only the correctly folded protein form of MFAP5 is detected, suggesting some mechanism which prevents the secretion of the truncated form. This is consistent with our secretion assay results, which demonstrated a 50-80% loss of MFAP2 secretion when the *O*-fucosylation site is mutated (T105A) or when *POFUT3/4* are knocked out. Interestingly, while MFAP5 secretion is reduced in *POFUT3/4* DKO HEK293 cells, this loss is not as severe as either MFAP2 or the N-terminal EMI control (**Fig 5i,k**). Furthermore, the MFAP5 alanine mutant (T92) secretion assay showed no significant change in secretion despite blockage of the *O*-fucosylation site **(Fig. 5c,d**). This may be due to the presence of a previously reported *N*-glycan with core fucose site (N79) in close proximity to the MFAP5 MBD^43^. This would also explain why secretion is only reduced to the same level as the EMI domain when GDP-fucose synthesis is entirely suspended in the *FX* KO cells (**Fig. 5j,l**). As POFUT3/4 modify only correctly folded proteins, the reported loss of MFAP5 secretion in patients may be due to these enzymes’ role in protein folding quality control. Future work should thus investigate the role of *O*-fucosylation through mouse *Pofut3/Pofut4* knockouts, examining for vascular defects or disturbance to aortic structure and function.

Structural comparison algorithms such as DALI have long been used to investigate protein function and infer evolutionary relationships^44^. However, these technologies suffer from long computing times that has heavily limited their potential applications^45,46^. Foldseek^47^ utilises an optimised 3Di algorithm to improve computing speed, and can leverage the entire AlphaFold predicted protein structure database^19^ to rapidly compare a query protein against entire proteomes. Here, we have demonstrated that iterative Foldseek analysis accurately identified several novel *O*-fucosylation substrates with the input of just one predicted protein structure, and we have validated these predictions in vivo using human tissue samples. These data suggest that there are likely many other protein domain families that only share structural homology, but not primary sequence homology, and yet participate in essential biochemical interactions. Using structural homology to screen putative enzyme-substrate interactions would be highly beneficial in protein interactome studies and for drug discovery^48^. Future work is required to explore applications of this methodology for other protein domain-specific interactions and enzyme-substrate relationships, such as POGLUTs and protein kinases^49,50^.

In conclusion, using state-of-the-art structural homology analysis we identified a new class of POFUT3/4 substrates that we referred to as EMI-like domains, which includes the MBDs of MFAP2 and MFAP5. For the first time, we observed a substrate preference between POFUT3 and POFUT4: *O*-fucosylation of MFAP2/MFAP5 is predominantly mediated by POFUT3 and is necessary for efficient secretion. All of these modified regions are disulfide-rich, resulting in an anti-parallel twisted beta sheet structure with high structural conservation despite lacking primary sequence homology. Our results contribute to the growing evidence that *O*-fucosylation is involved in a wide range of physiological processes and is required for a non-canonical protein folding quality control pathway, in which the transfer of *O*-fucose may act as a signal permitting release from the ER.

## Materials and Methods

### Foldseek iterative structural homology search

The predicted structure of human MMRN1-EMI domain (HUMAN_MMRN1: 207-282), a known POFUT3/POFUT4 substrate, was generated using Colabfold^21^. Templating from the Protein Data Bank was allowed and relaxation was not used. This structure was then searched in Foldseek^47^, to identify homologous human protein domains within the AlphaFold/UniProt50 (version 4) database. A taxonomic filter was used to restrict the search to *Homo sapiens* and search mode was set to 3Di/AA. All high scoring protein domains identified from this initial search were then searched individually in Foldseek. This process was repeated iteratively two more times. The probability, sequence identity and e-value scores were recorded at each stage to generate a protein homology network, and these raw values are available in Supplementary Table 1. Structural alignment to compare the predicted structures of MMRN1 EMI-domain and the MFAP2/MFAP5 matrix binding domain was performed in ChimeraX using the Matchmaker algorithm with default settings.

### AlphaFold2-multimer Interaction Analysis

An AlphaFold2-multimer interaction screen was performed using Colabfold^21^ to estimate the binding of human fucosyltransferase enzymes for the MBD of either MFAP2 (MFAP2_HUMAN: 97-143), or MFAP5, (MFAP5_HUMAN: 84-130). The residue ranges used for each FUT in this analysis were as follows: FUT1_HUMAN: 78–365; FUT2_HUMAN: 61–343; FUT3_HUMAN: 60–361; FUT4_HUMAN: 187–530; FUT5_HUMAN: 75–374; FUT6_HUMAN: 60–359; FUT7_HUMAN: 46–342; FUT8_HUMAN: 105–575; FUT9_HUMAN: 62–359; OFUT1_HUMAN: 28–388; OFUT2_HUMAN: 41–429; OFUT3_HUMAN: 81-479 and OFUT4_HUMAN 73-492. Templating from the Protein Data Bank was allowed and relaxation was not used.

### Patient Adipose Biopsy Collection

An adipose tissue biopsy was performed in the morning following a 12-hour overnight fast according to the technique of Bergstrom and previously described^51^. Briefly, this technique involves cleansing the skin on the abdomen lateral to the umbilicus with chlorhexidine solution, anaesthesia is administered (5 ml of Xylocaine 2%, no adrenaline). A 0.75 cm incision is made in the skin with a scalpel and a 5 mm Bergstrom needle inserted to collect approximately 200 mg of adipose tissue with suction. Two to three passes will be used to obtain the adipose tissue. The sample is blotted, and snap frozen as quickly as possible on dry ice. Upon completion of the biopsy, pressure is applied, and the incision closed with a sterile bandage, and a sterile dressing applied. The adipose tissue samples collected for this analysis were from three participants undertaking a randomized controlled trial. Ethics approval was obtained from the Human Research Ethics Committee of the University of Adelaide (H-2022-199). The study procedures, purpose, and known potential risks were explained to each participant according to the principles of the Declaration of Helsinki, and all participants will provide written informed consent before participation.

### Preparation of human adipose tissue for LC-MS/MS analysis

For analysis of in vivo *O*-fucosylation of MFAP2 T105 and MFAP5 T92, frozen human adipose tissue (~200 mg) was placed in 1 ml room temperature extraction buffer (1× cOmplete protease inhibitor EDTA-free (Roche), 4% SDS, 100 mM NaCl, 20 mM NaPO_4_). Homogenised tissue lysate was heated to 65°C for 10 min with mixing at 1,000 rpm, then sonicated for 10 min using a QSonica Q800R2 for 10 min total sonication time (30 s ON, 30 s OFF, 30% amplitude, water temperature 18 °C). The lysate was clarified by centrifugation at 18,000*g* for 10 min at 18 °C. The lipid layer formed at the top of the sample was removed by aspiration slowly. A portion of the lysate (50 μL) was transferred to a new tube and lipid removed by through chloroform-methanol protein precipitation^52^. Proteins were resuspended in 100 μl of resuspension buffer (4% SDC, 100 mM Tris-HCl pH 8.0) and vortexed until fully dissolved.

Protein concentration was measured using a BCA assay according to manufacturer’s instructions (Thermo Fisher Scientific, 23225). For protein digestion, 20 μg of lysate protein was reduced and alkylated by addition of chloroacetamide (CAA) and triscarboxyethylphosphine (TCEP) to final concentrations of 10 mM and 40 mM, respectively, and subsequently heated to 95 °C for 10 min with mixing at 1,000 rpm. Water was added to achieve a final SDC concentration of 1% and samples were allowed to reach room temperature before the addition of 1 μg of trypsin (Sigma-Aldrich) and lysC (Wako) for digestion (16 h, 37 °C). Peptide cleanup was performed using SDB-RPS StageTips^53^.

### LC-MS/MS analysis of *O*-fucosylated peptides in human adipose tissue using EAD Fragmentation

From each digest, 500 ng of peptides were loaded onto each Evotip Pure tip (EvoSep) after dilution with 0.1% formic acid (FA) in water according to the manufacturer’s instructions. Loaded Evotips were mounted on an EvoSep-1 HPLC coupled to a 150 μm × 8 cm C18 column via an Optiflow source with the column oven at 40 °C to a 7600 ZenoTOF instrument (Sciex). Samples were separated using the 60 samples per day (SPD) LC method and analysed using parallel reaction monitoring (PRM) targeting the MFAP2 peptide containing T105 (EEQYPC[Carbamidomethyl]T[Fuc]R *O*-fucosylated: precursor m/z = 614.7612, z = +2, product ion m/z = 533.2500; unmodified: *m/z* = 541.7322, z= +2, product ion m/z = 533.2500), or the MFAP5 peptide containing T92 (FTC[Carbamidomethyl]T[Fuc]R *O*-fucosylated: *O*-fucosylated: m/z *=* 415.6893, z= +2, product ion m/z = 537.2450; unmodified: *m/z* = 342.6603, z= +2, product ion m/z = 537.2450). EAD and collision induced dissociation parameters were optimised for each peptide ion^5^. EAD fragmentation spectra were averaged across the chromatographic peak and submitted to Byonic (Protein Metrics) for identification. The full human proteome (UniProt accession number UP000005640, downloaded on 14/02/24) was used with an FDR of 2% using a target-decoy-based strategy for protein and peptide identification. MS1 and MS2 mass tolerance was set to 4 ppm and 20 ppm, respectively. Trypsin was set as the digestion enzyme with a maximum of two missed cleavages. Carbamidomethylation of cysteine was set as a fixed modification and *O*-Fuc (1) was set as a rare variable modification.

### LC-MS/MS analysis of *O*-fucosylated peptides by PRM and DDA

From each digest, 500 ng of peptides were injected onto a NeoVanquish UHPLC coupled to a 75 μm × 50 cm C18 column via a nanosprayFLEX source with the column oven at 60 °C to a Thermo Scientific Eclipse or Exploris mass spectrometer. Samples were separated using a 25 min gradient and analysed using either data-dependent acquisition (DDA) with a 1.5 s cycle time, or parallel reaction monitoring (PRM) as per the adipose tissue analysis described above. DDA data were searched using MaxQuant with the full human proteome (UniProt) and a protein and peptide-level identification FDR of 1% was applied. Trypsin was set as the digestion enzyme with a maximum of two missed cleavages. Carbamidomethylation of cysteine was set as a fixed modification and *O*-Fuc(1) was set as a rare variable modification.

For glycoproteomic site mapping of MFAP2/MFAP5 proteins from HEK293T WT, *POFUT3* KO, *POFUT4* KO, *POFUT3/4* KO, *FX* KO cells or products from enzymatic assays, purified proteins (100 μl of elution from Ni-NTA purification), 300 μl of conditioned culture media or enzymatic assay products were precipitated using 3× volumes of cold acetone at −20 °C overnight, followed by centrifuging at 18,213g for 12 min at 4 °C. Pellets were denatured and reduced with 30 μl of reduction buffer (8 M urea, 0.4 mM ammonium bicarbonate and 10 mM TCEP), incubated at 60 °C for 10 min and cooled down to room temperature. 15 μl of 100 mM iodoacetamide was added, and samples were incubated in the dark for 40 min at room temperature, followed by diluting with 135 μl of water. Digestion was performed with 1 μg of trypsin (Thermo Fisher Scientific, 90057) for MFAP2 and N-terminal EMI samples or 1 μg of lysC (Thermo Fisher Scientific, 90307) for MFAP5 samples overnight at 37 °C. Samples were acidified with formic acid (FA) to 0.5%, sonicated for 20 min and desalted with a C18 Ziptip (Millipore, ZTC18S960). From each digest, ~10 ng of peptides was injected onto an EASY-nLC 1200 System (Thermo Fisher Scientific) coupled to an Acclaim PepMap-100 75 μm × 2 cm nanoViper C18 pre-column, an EasySpray PepMap RSLC 50 μm × 15 cm C18 analytical column and a Q Exactive Plus mass spectrometer (Thermo Fisher Scientific) with Tune 2.12 (Build 3134). Peptides were gradient eluted at a constant flow rate of 300 nl per min using a 90-min gradient and a 120-min instrument method. The gradient profile was as follows: (min: % Solvent B (80%ACN/0.1% FA)) 0:0, 90:50, 93:98, 96:98, 99:2, 102:2, 105:98, 108:98, 111:2, 114:2, 117:98, 120:98. Data was collected with data-dependent acquisition (DDA). MS1 scans were acquired with a resolution of 70,000, an AGC target of 1 × 10^6^ and a mass range from 400 m/z to 2,000 m/z. Dynamic exclusion was enabled for 6 s. MS2 scans were acquired in Orbitrap with a resolution of 17,500, an isolation window of 1.2 m/z and an AGC target of 1 × 10^5^. Higher-energy collisional dissociation (HCD) fragmentation was set with a fixed collision energy of 27%. Data analysis was performed with Byonic software (Protein Metrics, 4.1.10). Search parameters include fully specific cleavage specificity at the C-terminal of R and K for trypsin or K for lysC with two missed cleavages allowed. Mass tolerance was set at 10 ppm for precursors and 20 ppm for fragments. Cysteine carbamidomethylation was set as fixed modification. Methionine oxidation (common1) and N-terminal acetylation (rare1) were set as variable modifications. Glycans were set as varaiable modifications (common1) using a customed *O*-glycan search space including Fuc(1), HexNAc(1)Fuc(1), HexNAc(1)Hex(1)Fuc(1), HexNAc(1)Hex(1)Fuc(1)NeuAc(1), Hex(1)Fuc(1), Hex(1), Hex(1)Pent(1), Hex(1)Pent(2) and HexNAc(1). To make the EICs for a given peptide, the ions of each glycoform were extracted using calculated m/z with a mass tolerance of ±0.005, and then overlaid to compare the relative ion intensity. EICs were smoothed using a Gauss algorithm. All EICs were manually extracted and anaylsed using Xcalibur (Thermo Fisher Scientific, version 4.0.27.19).

### Cell culture

CRISPR-Cas9 HEK293T KOs of *POFUT3, POFUT4* and *FX* were generated in Hao et al^5^. HEK293T WT, *POFUT3* KO, *POFUT4* KO, *POFUT3/4* KO and *FX* KO cells were cultured in Dulbecco’s Modified Eagle Medium (DMEM, GE Healthcare Life Sciences) and supplemented with 10% bovine calf serum (BCS, VWR 1058-358), 100 U ml^-1^ penicillin and 100 μg ml^-1^ streptomycin. Adapted suspension culture of HEK293T *FX* KO cells were cultured in Freestyle medium (Gibco). Cells were incubated at 37 °C with 5% CO_2_. HEK293T cells were purchased from the American Type Culture Collection (ATCC).

### Plasmids and mutagenesis

The mammalian expression plasmids including pcDNA3.1-hMFAP2 WT-FLAG, pcDNA3.1-hMFAP2 T105A-FLAG, pcDNA3.1-hMFAP5 WT-FLAG, pcDNA3.1-hMFAP5 T92A-FLAG, pcDNA3.1-hMFAP2-MycHis and pcDNA3.1-hMFAP5-MycHis were synthesised by Genescript. pcDNA4-hMMRN1 N-terminal EMI-MycHis, pcDNA4-full-length hPOFUT3-MycHis and pcDNA4-full-length hPOFUT4-MycHis plasmids were described previously^5^. pGEn2-GFP, pGEn2-GFP-hPOFUT3 (FUT10) and pGEn2-GFP-hPOFUT4 (FUT11) plasmids were described previously and were generously provided by Kelly Moremen at the University of Georgia^25^.

### Adaption of HEK293T *FX* KO adherent cells to serum-free suspension

The protocol was adapted from Lomba et al^26^. Briefly, adherent HEK293T *FX* KO cells grown in 10 cm dishes with DMEM with 10% BCS and 1% penicillin/streptomycin (adhesion media) were adapted to serum-free condition by sequential passages in gradually increasing proportions of Freestyle medium (adhesion media: serum-free medium ratios of 0%, 25%, 50%, 75%, 100%) containing 1% penicillin/streptomycin. Two passages were performed per condition with cells reaching at least 70% confluency at the time of passaging. Once adapted to serum-free condition, cells were transferred to a 125 ml Erlenmeyer flask containing 20 ml of serum-free media and maintained in an orbital shaker at 150 rpm, 37 °C with 5% CO_2_.

### Co-immunoprecipitation of POFUT3/4 with substrate proteins

Recombinant POFUT3 and POFUT4 (GFP-tagged, no transmembrane domain) were expressed in HEK293T cells alongside FLAG-tagged substrate proteins (MFAP2, MFAP5). Cells were seeded in 10-cm dishes in 10 ml of DMEM overnight to achieve 70% confluency. Cells were then transiently transfected using Lipofectamine 3000 (Thermo Fisher Scientific) with 10.5 μg of substrate plasmid (either pcDNA3.1-hMFAP2 WT-FLAG, pcDNA3.1-hMFAP5 WT-FLAG, or pcDNA3.1 EV) alongside 10.5 μg of enzyme plasmid (pGEn2-GFP-hPOFUT3, pGEn2-GFP-hPOFUT4, or pGEn2-GFP EV). After two days, cells were lysed and co-immunoprecipitated with Pierce protein G magnetic beads (Thermo Scientific) and anti-FLAG M2 antibody (Sigma Aldrich) as reported previously in Hao et al^5^. Eluates were digested with trypsin and peptides clean-up as previously described^53^.

### Production of MFAP2/MFAP5 proteins used for mass spectral analysis from *POFUT3/4* and *FX* knockout cells

Generation of all CRISPR-Cas9 HEK293T KOs, including *POFUT3, POFUT4, POFUT3*/*4* DKOs and *FX* KOs, was described previously in Hao et al^5^. HEK293T WT, *POFUT3* KO, *POFUT4* KO, *FX* KO and *POFUT3*/*4* DKO cells were seeded in 10-cm dishes and cultured overnight to reach a confluency of 70%. Each plate of cells were transiently transfected with 5 μg of pcDNA3.1-hMFAP2-MycHis or 10 μg of pcDNA3.1-hMFAP5-MycHis in 8 ml of Opti-MEM (Thermo Fisher Scientific) using PEI (6 μg PEI per 1 μg DNA). Two days later, media from 10 plates of cells were combined. Proteins were purified using Ni-NTA agarose (Qiagen) and eluted with 250 mM imidazole in Tris-buffered saline (TBS), pH 7.5. Purified proteins were stored at −20 °C until use for glycoproteomic mass spectrometry analysis as described above. For POFUT3/4 rescue assays of MFAP2, *POFUT3/4* DKO cells (1 × 10^6^ cells per well) were seeded in six-well plates and transfected with 2 μg of pcDNA3.1-hMFAP2-MycHis with 0.2 μg of pcDNA4-full-length hPOFUT3-MycHis, pcDNA4-full-length hPOFUT4-MycHis or empty vector in 1.2 ml of Opti-MEM using PEI (6 μg PEI per 1 μg DNA). Cells were cultured for 2 days and media were collected and digested for mass spectrometry analysis as described above. For POFUT3/4 rescue assays of MFAP5, *POFUT3/4* DKO cells were seeded in 10-cm dishes and transfected with 8 μg of pcDNA3.1-hMFAP5-MycHis with 0.8 μg of pcDNA4-full-length hPOFUT3-MycHis, pcDNA4-full-length hPOFUT4-MycHis or empty vector in 8 ml of Opti-MEM using PEI (6 μg PEI per 1 μg DNA). Cells were cultured for 2 days. Proteins were purified from the media of 5-6 plates using Ni-NTA agarose (Qiagen) and use for glycoproteomic mass spectrometry analysis as described above.

### *In vitro* enzymatic assays using purified MFAP2/MFAP5

MFAP2 enzymatic assays were performed using purified, non-fucosylated MFAP2 mixed with recombinant GFP-POFUT3/4 (lacking transmembrane domain). Recombinant GFP, GFP-POFUT3 and GFP-POFUT4 were purified from the media of HEK293F cells transfected with pGEn2-GFP, pGEn2-GFP-hPOFUT3 and pGEn2-GFP-hPOFUT4 plasmids, respectively^25^. Transfections were performed with 4 μg DNA per ml and 9 μg PEI per ml. Non-fucosylated MFAP2 was purified from HEK293T *FX* KO cells adapted to serum-free suspension culture as described above^26^. Transfections were performed with pcDNA3.1-hMFAP2-MycHis (4 μg DNA per ml) and PEI (9 μg per ml). Non-fucosylated N-terminal EMI (positive control) was purified from HEK293F cells that incubated with 200 μM 6-alkynyl fucose as described previously^5^. Transfections were performed with pcDNA4-hMMRN1 N-terminal EMI-MycHis (4 μg DNA per ml) and PEI (9 μg per ml). Cells were cultured for 3 days. Proteins were purified from the media using Ni-NTA agarose (Qiagen). The purity of proteins was verified by Coomassie blue staining. For acquiring the time-dependent activity of the enzymes, 0.1 μM recombinant GFP–POFUT3, GFP– POFUT4 or GFP (negative control) was incubated in 150 μl reaction mixtures (for MFAP2) or 50 reaction mixtures (for N-terminal EMI) containing 100 μM GDP-fucose, 300 μM MnCl and 0.5 μM purified non-fucosylated MFAP2 or N-terminal EMI in 50 mM HEPES, pH 7. Reactions were incubated at 37 °C for indicated times and stopped by adding 10× volume of cold acetone. Reaction products were analysed with nano LC-MS/MS as described earlier.

For MFAP5 enzymatic assays, non-fucosylated MFAP5 MBD (residues 84-129) was generated synthetically with heated automated Fmoc-SPPS performed on a Biotage SYRO I automated synthesizer, as described in **Supplementary Methods 1**. To acquire the time-dependent activity of the enzymes, 0.5 μM purified non-fucosylated MFAP5 MBD was incubated in 20 μl reaction mixture containing 100 μM GDP-fucose, 300 μM MnCl and 50 mM HEPES, pH 7, alongside either 0.1 μM recombinant GFP-POFUT3, GFP-POFUT4, or no enzyme. Reactions were incubated at 37 °C for indicated times. Reaction products were analysed with nano LC-MS/MS as described earlier.

For bacterially expressed MFAP2 enzymatic assays, non-fucosylated MFAP2 was expressed as described below. To acquire the time-dependent activity of the enzymes, 0.5 μM purified non-fucosylated MFAP2 was incubated in 20 μl reaction mixture containing 100 μM GDP-fucose, 300 μM MnCl and 50 mM HEPES, pH 7, alongside either 0.1 μM recombinant GFP-POFUT3, GFP-POFUT4, or no enzyme. Reactions were incubated at 37 °C for indicated times. Reaction products were analysed with nano LC-MS/MS as described earlier.

### Expression of MFAP2 in Bacteria

SHuffle T7 Express Competent *E. coli* (NEB, Cat#C3029J) were transformed by the heat shock method with pET-32a-human MFAP2 plasmid. 50 µL of *E. coli* was thawed on ice for 10 minutes before 2.5 µL of purified plasmid DNA (200 ng) was added to the E. coli transformation tube. The E. coli transformation tube was gently flicked five times to mix the *E. coli* and DNA. The transformation tube was incubated on ice for 30 min followed by 30 s at 42°C in a pre-warmed water bath. The reaction tube was moved directly from the water bath to ice for a 5 min incubation. 950 µL of room temperature SOC media was added and incubated at 30°C for 1 hour with shaking at 250 rpm. The transformation was plated on LB agar containing 100 μg/mL ampicillin antibiotic and incubated at 30°C for 36 h.

One colony was selected and transferred into 10 mL of LB broth containing 100 μg/mL ampicillin in a 50mL falcon tube and grown overnight at 30°C with 200 rpm shaking. The bacteria were diluted 100-fold into 500 mL of pre-warmed LB broth containing 100 μg/mL ampicillin in 2.5 L flask and grown at 30°C with 200 rpm shaking to an OD_600nm_ of 0.5-0.7. The culture was then cooled to 16°C, and 100 mM isopropyl ß-d-1 thiogalactopyranoside (IPTG) stock in water was added to a final concentration of 0.1 mM IPTG and incubated for 24 h at 16oC for protein expression.

The bacteria were harvested by centrifugation at 8,000 g for 10 min at 4°C. The supernatant was discarded and the pellet frozen at −30°C. The pellet was thawed on ice for 10 min then resuspended in 10 mL lysis buffer (20 mM sodium phosphate, 500 mM sodium chloride, 1X cOmplete, EDTA-free Protease Inhibitor, 0.2 mg/mL Lysozyme (Sigma Cat#L6876-1G), 20 µg/mL DNase (Sigma Cat#DN25-10MG), 1 mM magnesium chloride, pH 7.4). Samples were maintained on ice in 2 mL tubes and sonicated (QSonica Q800R2) for 10 minutes total sonicating time at 30% amplitude with 30s on and 30s off at 4°C. The lysate was centrifuged at 50,000 g for 30 min at 4°C. The supernatant and pellet were collected, but all MFAP2-6xHis tagged protein was in the pellet (inclusion bodies). The pellet was resuspended by vortexing in inclusion body lysis buffer (6 M Guanidine-HCl, 20 mM sodium phosphate, 500 mM NaCl, 5 mM imidazole, pH 7.4) at room temperature and incubated for 1 h. The solution was then centrifuged at 50,000 g for 30 min at 18°C and the supernatant collected. Equilibrated Ni-EXCEL beads (1 mL slurry, Cytiva Cat#17371201) were added to the inclusion body supernatant and incubated for 30 min at room temperature. The beads were then pelleted by centrifugation at 2,000 g for 2 min (slow deceleration) at RT, the supernatant removed and the beads resuspended in 2 mL of inclusion body lysis buffer. The bead slurry was loaded into a 10 mL gravity flow column (Thermo) and washed three times with 10 mL of inclusion body lysis buffer at RT. The protein was refolded with 10 ml each of inclusion body lysis buffer with decreasing concentrations of guanidine-HCl (5 M, 4 M, 3 M, 2 M, 1 M) at RT. The column was then washed three times with 10 ml wash buffer (20 mM sodium phosphate, 500 mM NaCl, 20 mM imidazole, pH 7.4) at RT. MFAP2 was eluted with 5 mL of elution buffer (20 mM sodium phosphate, 500 mM NaCl, 500 mM imidazole, pH 7.4) at RT and stored at 4°C with the addition of 6% sodium azide to 0.02% final concentration.

### Western-blot based secretion assays

To analyse the secretion of MFAP2 (WT/T105A) and MFAP5 (WT/T92A) expressed in HEK293T WT cells, cells were seeded in 6 well plates (0.4 × 10^6^ cells per well) and incubated overnight. Cell media was changed to Opti-MEM once 80% confluency was achieved, and cells were then transfected with Lipofectamine 3000. Each well was transfected with 2 μg of either pcDNA3.1-hMFAP2 WT-FLAG, pcDNA3.1-hMFAP5 WT-FLAG, pcDNA3.1-hMFAP2 T105A-FLAG, pcDNA3.1-hMFAP5 T92A-FLAG, or empty vector. 0.5 μg of IgG plasmid was co-transfected as a secretion control. Media samples were collected after 48 hours of incubation and filtered with 0.45-μm filters (Millipore). Media (200 μl) from each well was precipitated with chloroform-methanol precipitation. The resulting pellet was then resuspended in 50 μL of 1x LDS buffer with TCEP (10 mM) and was immediately heated to 65 °C for 10 min. Samples were loaded onto a 4-12% Bolt Bis-Tris Plus Protein Gel (Invitrogen) with MES buffer (200 V, 25 min) and transferred to a nitrocellulose membrane. Membranes were then blocked with 5% skim-milk for 90 min at room temperature. For MFAP2-FLAG secretion assays, membranes were incubated with mouse anti-FLAG antibody (Sigma Aldrich, 1:1,000) overnight at 4 °C. Membranes were then incubated with 680-conjugated goat anti-human IgG antibody (LI-COR, 1:15,000) and 800-conjugated goat anti-mouse IgG antibody (LI-COR, 1:15,000) for 1 h at room temperature. For MFAP5-FLAG secretion assays, membranes were incubated with rabbit anti-human MFAP5 primary antibody (Proteintech Group, 1:1,000) for 16 h at 4 °C. Membranes were then incubated with 680-conjugated goat anti-human IgG antibody (LI-COR, 1:15,000) and 800-conjugated goat anti-rabbit IgG antibody (Rockland Immunochemicals, 1:15,000) for 1 h at room temperature. To analyse the secretion of MFAP2 and MFAP5 expressed in HEK293T WT, *FX* KO and *POFUT3*/*4* DKO cells, cells were seeded in 6 well plates (1 × 10^6^ cells per well) and incubated overnight to reach a confluency of 80%. Cell media was changed to 1 ml of Opti-MEM. For each well, cells were transfected with 0.5 μg of pcDNA3.1-hMFAP2-MycHis, 5 μg of pcDNA3.1-hMFAP5-MycHis, 0.5 μg of pcDNA4-hMMRN1 N-terminal EMI-MycHis (positive control) or empty vector, together with 0.1 μg of IgG plasmid using PEI (6 μg PEI per 1 μg DNA). Media samples were collected after 48 hours of incubation. Media (100 μl) were precipitated with 500 μl of cold acetone overnight at −20 °C and centrifuged at 18,213*g* for 12 min to obtain pellets. Pellets were resuspended in 12 μl of 2× reducing buffer (4% SDS, 200 mM 2-mercaptoethanol, 20% glycerol in 100 mM Tris/HCl, pH 6.8) by sonication for 10 min and boiled at 105 °C for 8 min. Samples were loaded on 4-20% SDS-PAGE gels (Bio-Rad) and transferred to a nitrocellulose membrane. Membranes were blocked with 5% non-fat milk (Bio-Rad) for 30 min at room temperature followed by incubating with anti-Myc antibody (Invitrogen, clone 9E10, 1:2,500) overnight at 4 °C. Membranes were then incubated with IDRye 800-conjugated goat anti-mouse IgG antibody (LI-COR, 1:2,500) and IDRye 680-conjugated goat anti-human IgG antibody (LI-COR, 1:2,500) for 1 h at room temperature. All western blot bands were visualised and quantified using an Odyssey system (LI-COR) with LI-COR Image Studio (version 5.2.5).

## Acknowledgements

We thank all the current and past members of the Haltiwanger laboratory for technical advice and critical comments on this paper. We thank SydneyMS for providing instrumentation used in this study. This work was supported by National Institutes of Health grant R35GM148433 awarded to R.S.H., an ARC Laureate Fellowship (FL250100011) awarded to R.J.P. and National Health and Medical Research Council grant (GNT2030089) awarded to M.L.

## Notes

### Competing Interest Statement

The authors have declared no competing interest.

### Summary of Updates

A typographical error in an author name was corrected.

## References

1. Hao, H., Eberand, B. M., Larance, M. & Haltiwanger, R. S. Protein O-Fucosyltransferases: Biological Functions and Molecular Mechanisms in Mammals. Molecules. 30 (2025).

2. Holdener, B. C. & Haltiwanger, R. S. Protein O-fucosylation: structure and function. Curr. Opin. Struct. Biol. 56, 78–86 (2019).

3. Rampal, R., Arboleda-Velasquez, J. F., Nita-Lazar, A., Kosik, K. S. & Haltiwanger, R. S. Highly conserved O-fucose sites have distinct effects on Notch1 function. J. Biol. Chem. 280, 32133–32140 (2005).

4. Neupane, S. et al. O-fucosylation of thrombospondin type 1 repeats is essential for ECM remodeling and signaling during bone development. Matrix Biol. 107, 77–96 (2022).

5. Hao, H. L. et al. FUT10 and FUT11 are protein O-fucosyltransferases that modify protein EMI domains. Nat. Chem. Biol. 21 (2025).

6. Luo, Y. & Haltiwanger, R. S. O-fucosylation of notch occurs in the endoplasmic reticulum. J. Biol. Chem. 280, 11289–11294 (2005).

7. Luo, Y., Koles, K., Vorndam, W., Haltiwanger, R. S. & Panin, V. M. Protein O-fucosyltransferase 2 adds O-fucose to thrombospondin type 1 repeats. J. Biol. Chem. 281, 9393–9399 (2006).

8. Li, Z. J. et al. Recognition of EGF-like domains by the Notch-modifying O-fucosyltransferase POFUT1. Nat. Chem. Biol. 13, 757–+ (2017).

9. Valero-González, J. et al. A proactive role of water molecules in acceptor recognition by protein O-fucosyltransferase 2. Nat. Chem. Biol. 12, 240–+ (2016).

10. Berardinelli, S. J. et al. O-fucosylation stabilizes the TSR3 motif in thrombospondin-1 by interacting with nearby amino acids and protecting a disulfide bond. J. Biol. Chem. 298 (2022).

11. Vasudevan, D., Takeuchi, H., Johar, S. S., Majerus, E. & Haltiwanger, R. S. Peters Plus Syndrome Mutations Disrupt a Noncanonical ER Quality-Control Mechanism. Curr. Biol. 25, 286–295 (2015).

12. Chabanais, J., Labrousse, F., Chaunavel, A., Germot, A. & Maftah, A. POFUT1 as a Promising Novel Biomarker of Colorectal Cancer. Cancers. 10 (2018).

13. Li, M. et al. Mutations in POFUT1, encoding protein O-fucosyltransferase 1, cause generalized Dowling-Degos disease. Am. J. Hum. Genet. 92, 895–903 (2013).

14. Shi, S. L. & Stanley, P. Protein O-fucosyltransferase 1 is an essential component of Notch signaling pathways. P. Natl. Acad. Sci. U.S.A. 100, 5234–5239 (2003).

15. Du, J. G. et al. O-fucosylation of thrombospondin type 1 repeats restricts epithelial to mesenchymal transition (EMT) and maintains epiblast pluripotency during mouse gastrulation. Dev. Biol. 346, 25–38 (2010).

16. Houlahan, C. B. et al. Analysis of the Healthy Platelet Proteome Identifies a New Form of Domain-Specific O-Fucosylation. Mol. Cell. Proteomics. 23 (2024).

17. Schneider, M., Al-Shareffi, E. & Haltiwanger, R. S. Biological functions of fucose in mammals. Glycobiology. 27, 601–618 (2017).

18. Zhang, Y. & Skolnick, J. TM-align: a protein structure alignment algorithm based on the TM-score. Nucleic Acids Res. 33, 2302–2309 (2005).

19. Fleming, J. et al. AlphaFold Protein Structure Database and 3D-Beacons: New Data and Capabilities. J. Mol. Biol. 437, 168967 (2025).

20. Combs, M. D. et al. Microfibril-associated Glycoprotein 2 (MAGP2) Loss of Function Has Pleiotropic Effects. J. Biol. Chem. 288, 28869–28880 (2013).

21. Mirdita, M. et al. ColabFold: making protein folding accessible to all. Nat. Methods. 19, 679–682 (2022).

22. Gilchrist, C. L. M., Mirdita, M. & Steinegger, M. Multiple protein structure alignment at scale with FoldMason. Science. 391, 485–488 (2026).

23. Doliana, R., Bot, S., Bonaldo, P. & Colombatti, A. EMI, a novel cysteine-rich domain of EMILINs and other extracellular proteins, interacts with the gC1q domains and participates in multimerization. FEBS Lett. 484, 164–168 (2000).

24. Kufareva, I. & Abagyan, R. Methods of protein structure comparison. Methods Mol. Biol. 857, 231–257 (2012).

25. Moremen, K. W. et al. Expression system for structural and functional studies of human glycosylation enzymes. Nat. Chem. Biol. 14, 156–162 (2018).

26. Lomba, A. L. O. et al. Serum-Free Suspension Adaptation of HEK-293T Cells: Basis for Large-Scale Biopharmaceutical Production. Braz. Arch. Biol. Technol. 64 (2021).

27. Lobstein, J. et al. SHuffle, a novel Escherichia coli protein expression strain capable of correctly folding disulfide bonded proteins in its cytoplasm. Microb. Cell Fact. 11, 56 (2012).

28. Ricketts, L. M., Dlugosz, M., Luther, K. B., Haltiwanger, R. S. & Majerus, E. M. -fucosylation is required for ADAMTS13 secretion. J. Biol. Chem. 282, 17014–17023 (2007).

29. Takeuchi, H. et al. -Glycosylation modulates the stability of epidermal growth factor-like repeats and thereby regulates Notch trafficking. J. Biol. Chem. 292, 15964–15973 (2017).

30. Sindelar, C. V., Hendsch, Z. S. & Tidor, B. Effects of salt bridges on protein structure and design. Protein Sci. 7, 1898–1914 (1998).

31. Consortium, G. T. The Genotype-Tissue Expression (GTEx) project. Nat. Genet. 45, 580–585 (2013).

32. Uhlen, M. et al. Proteomics. Tissue-based map of the human proteome. Science. 347, 1260419 (2015).

33. Wang, J. et al. Proteome Profiling Outperforms Transcriptome Profiling for Coexpression Based Gene Function Prediction. Mol. Cell. Proteomics. 16, 121–134 (2017).

34. Craft, C. S. et al. The extracellular matrix protein MAGP1 supports thermogenesis and protects against obesity and diabetes through regulation of TGF-beta. Diabetes. 63, 1920–1932 (2014).

35. Weinbaum, J. S. et al. Deficiency in microfibril-associated glycoprotein-1 leads to complex phenotypes in multiple organ systems. J. Biol. Chem. 283, 25533–25543 (2008).

36. Werneck, C. C. et al. Mice lacking the extracellular matrix protein MAGP1 display delayed thrombotic occlusion following vessel injury. Blood. 111, 4137–4144 (2008).

37. Broekelmann, T. J., Bodmer, N. K. & Mecham, R. P. Identification of the growth factor-binding sequence in the extracellular matrix protein MAGP-1. J. Biol. Chem. 295, 2687–2697 (2020).

38. Chen, E., Larson, J. D. & Ekker, S. C. Functional analysis of zebrafish microfibril-associated glycoprotein-1 (Magp1) in vivo reveals roles for microfibrils in vascular development and function. Blood. 107, 4364–4374 (2006).

39. Werneck, C. C. et al. Mice lacking the extracellular matrix protein MAGP1 display delayed thrombotic occlusion following vessel injury. Blood 111, 4137–4144 (2008).

40. Barbier, M. et al. MFAP5 loss-of-function mutations underscore the involvement of matrix alteration in the pathogenesis of familial thoracic aortic aneurysms and dissections. Am. J. Hum. Genet. 95, 736–743 (2014).

41. Ten Dijke, P. & Arthur, H. M. Extracellular control of TGFbeta signalling in vascular development and disease. Nat. Rev. Mol. Cell Biol. 8, 857–869 (2007).

42. Chen, Q. et al. Smad7 is required for the development and function of the heart. J. Biol. Chem. 284, 292–300 (2009).

43. Lih, T. M., Cho, K. C., Schnaubelt, M., Hu, Y. & Zhang, H. Integrated glycoproteomic characterization of clear cell renal cell carcinoma. Cell Rep. 42, 112409 (2023).

44. Holm, L. Dali server: structural unification of protein families. Nucleic Acids Res. 50, W210–W215 (2022).

45. Ma, J. & Wang, S. Algorithms, applications, and challenges of protein structure alignment. Adv. Protein Chem. Struct. Biol. 94, 121–175 (2014).

46. Hasegawa, H. & Holm, L. Advances and pitfalls of protein structural alignment. Curr. Opin. Struct. Biol. 19, 341–348 (2009).

47. van Kempen, M. et al. Fast and accurate protein structure search with Foldseek. Nat. Biotechnol. 42 (2024).

48. Miller, E. B. et al. Enabling structure-based drug discovery utilizing predicted models. Cell. 187, 521–525 (2024).

49. Li, Z. et al. Structural basis of Notch O-glucosylation and O-xylosylation by mammalian protein-O-glucosyltransferase 1 (POGLUT1). Nat. Commun. 8, 185 (2017).

50. Brinkworth, R. I., Breinl, R. A. & Kobe, B. Structural basis and prediction of substrate specificity in protein serine/threonine kinases. Proc. Natl. Acad. Sci. U.S.A. 100, 74–79 (2003).

51. Alderete, T. L. et al. A novel biopsy method to increase yield of subcutaneous abdominal adipose tissue. Int. J. Obes. (Lond). 39, 183–186 (2015).

52. Wessel, D. & Flugge, U. I. A method for the quantitative recovery of protein in dilute solution in the presence of detergents and lipids. Anal. Biochem. 138, 141–143 (1984).

53. Harney, D. J. et al. Proteomic Analysis of Human Plasma during Intermittent Fasting. J. Proteome Res. 18, 2228–2240 (2019).

54. Giansanti, P. et al. Mass spectrometry-based draft of the mouse proteome. Nat. Methods. 19, 803–811 (2022).

