## Supplementary Figures for "Iterative structural homology search identifies new substrates of the protein *O*-fucosyltransferases POFUT3 and POFUT4"

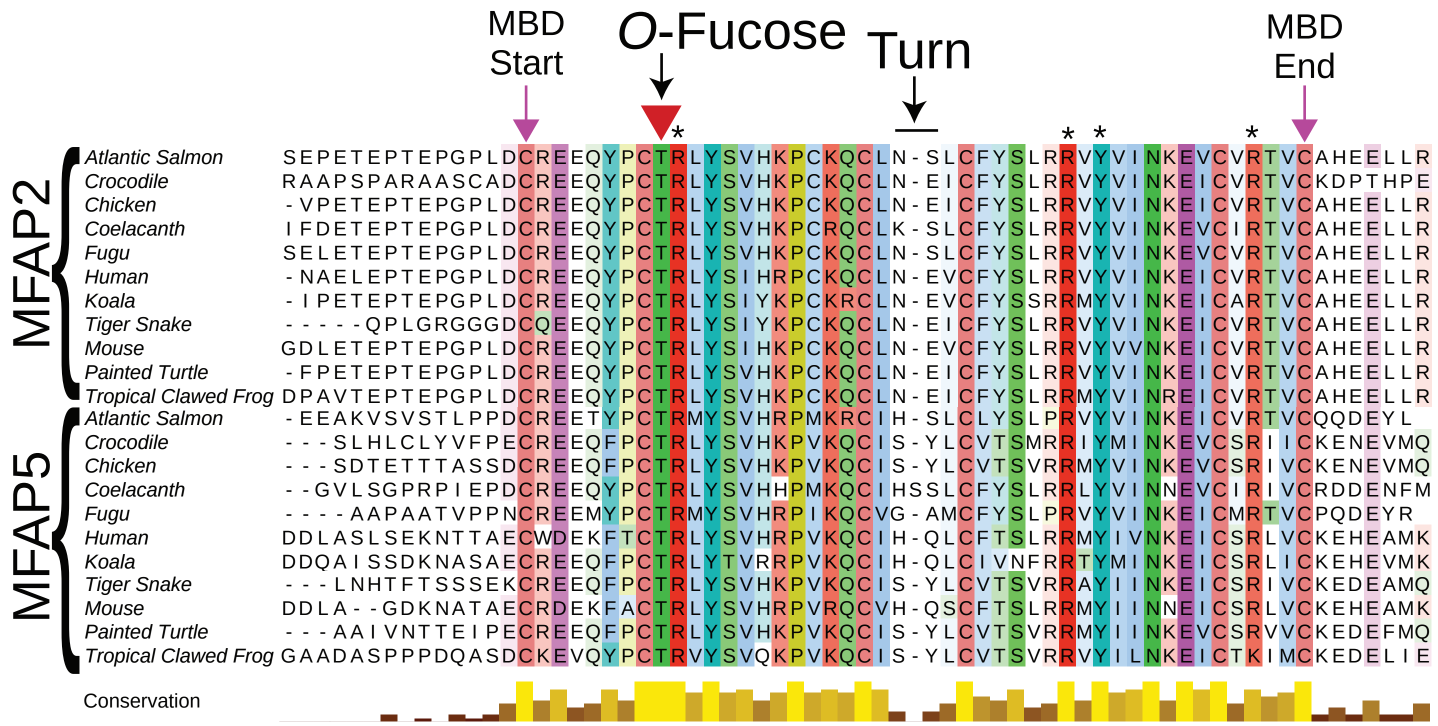


**Supplementary Fig. 1. Conservation of the MFAP2 and MFAP5 matrix binding domains**. Clustal Omega primary sequence alignment of the matrix binding domain of MFAP2 and MFAP5 across all vertebrate evolution, coloured by residue type. The site of *O*-fucosylation is shown by the red arrow and is fully conserved. Other residues which may be important for enzyme-substrate interactions due to their high conservation are also shown by asterisks. The flexible turn sequence is also shown to be the only significantly variable region of the domain.

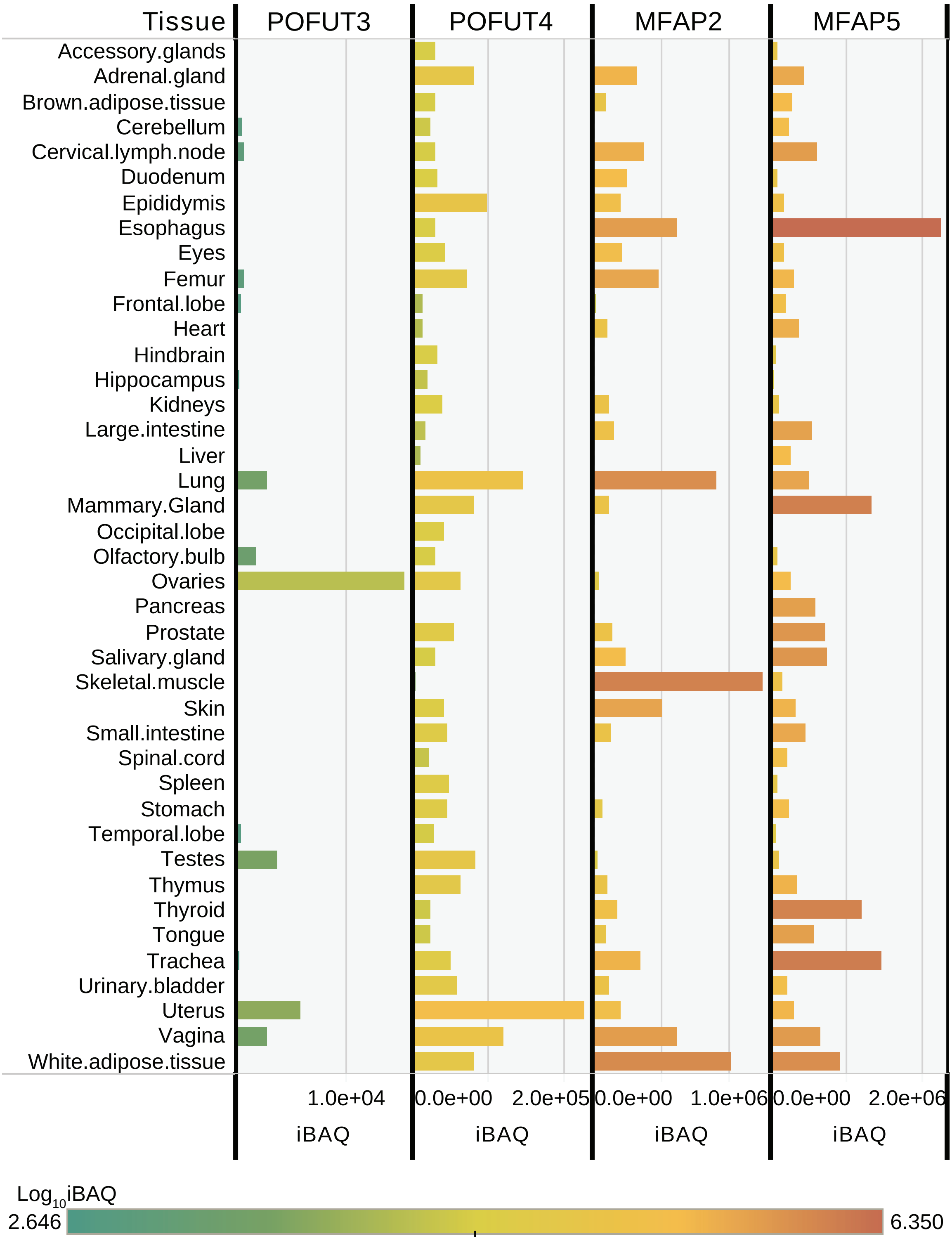


**Supplementary Fig. 2. Mouse tissue protein abundance profiles for key proteins of interest.** Normalised (iBAQ) abundance of POFUT3, POFUT4, MFAP2 and MFAP5 across murine tissue samples. Adapted from Giansanti *et al.*[54].

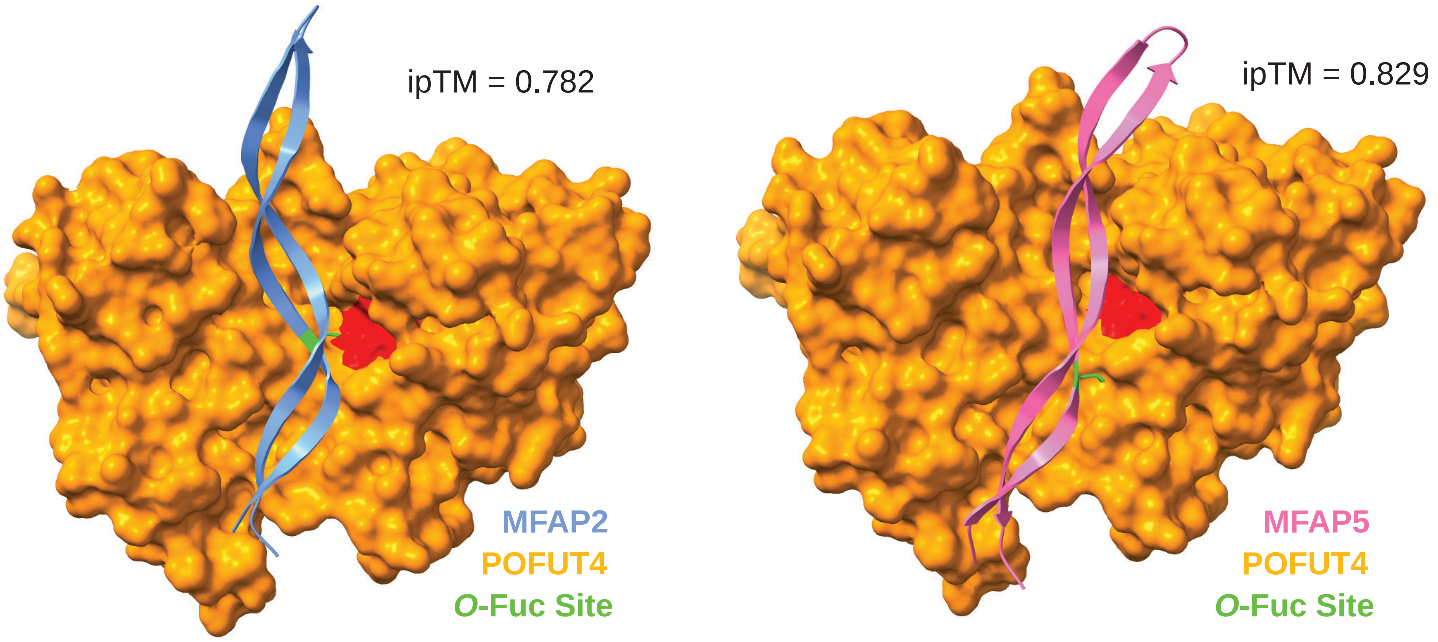


**Supplementary Fig. 3. POFUT4 interaction analysis using AlphaFold2**. AlphaFold2-multimer predicted structure of the interaction between human POFUT4 (orange, residues 73-492) and the matrix binding domain of human MFAP2 (left, residues 97-143) or MFAP5 (right, residues 84-130). The putative *O*-fucosylation site is highlighted in green, while the active site of the enzyme is highlighted in red. The interface predicted template modelling (ipTM) score is provided for each interaction.

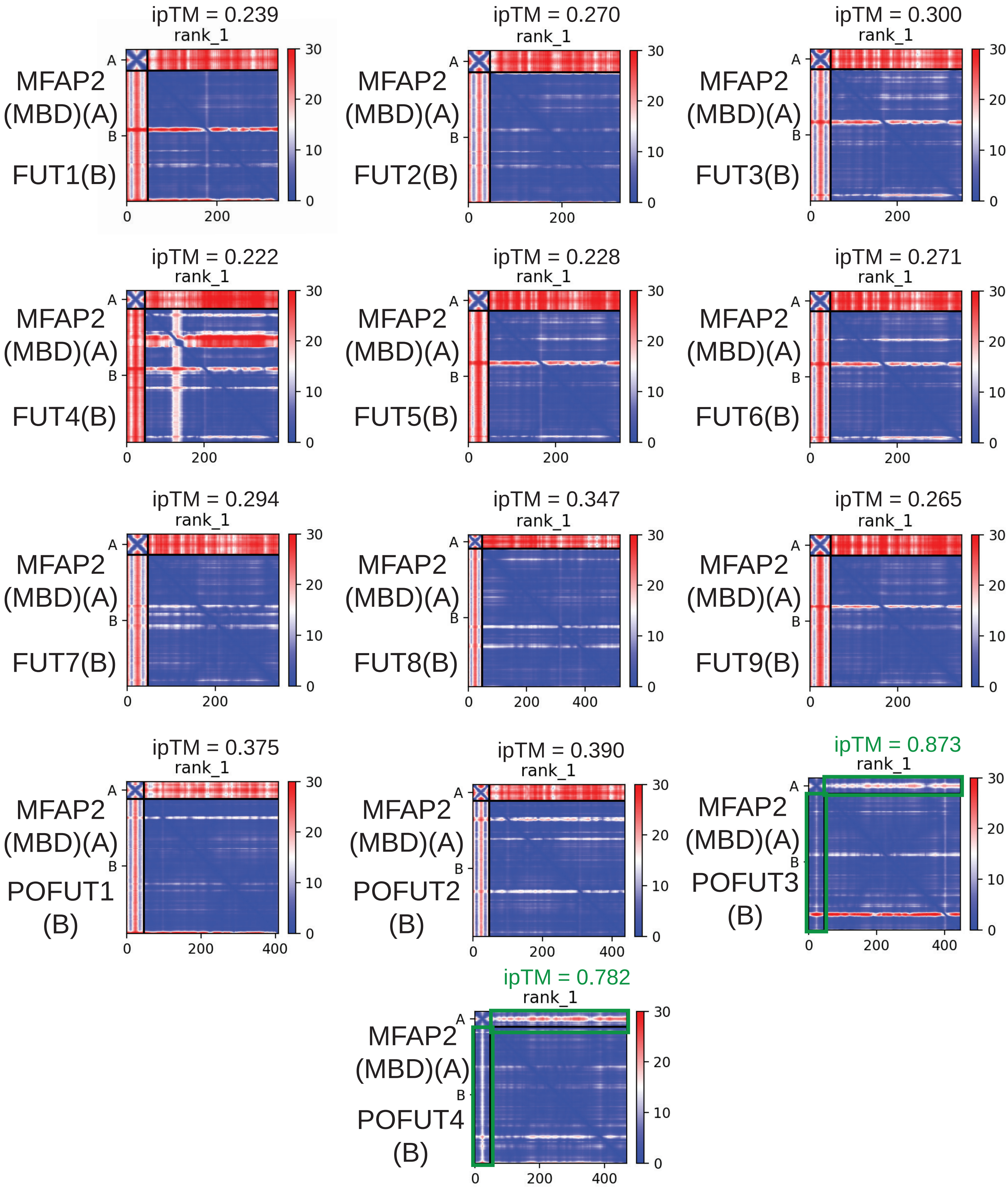


**Supplementary Fig. 4. FUT interaction screen with MFAP2 MBD.** PAE plots for the human MFAP2-fucosyltransferase AlphaFold2-multimer binding assay performed in Colabfold[21]. In each plot the MBD is molecule "A" and the fucosyltransferase is molecule "B". The top left and bottom right regions of each plot display the predicted aligned error (PAE) within either "A", or "B", respectively. The top right and bottom left regions of each plot display the PAE between "A" and "B". PAE is recorded in Angstroms, and lower values (blue) in the top right and bottom left regions of each plot indicate higher confidence for the interaction. These high confidence interactions are highlighted for POFUT3-MBD and POFUT4-MBD by green boxes. All other fucosyltransferases have very high error in these regions, indicating low confidence for their interaction. ipTM scores are provided for each plot, which is a quantitative measurement of the accuracy of the predicted interaction between the two molecules.

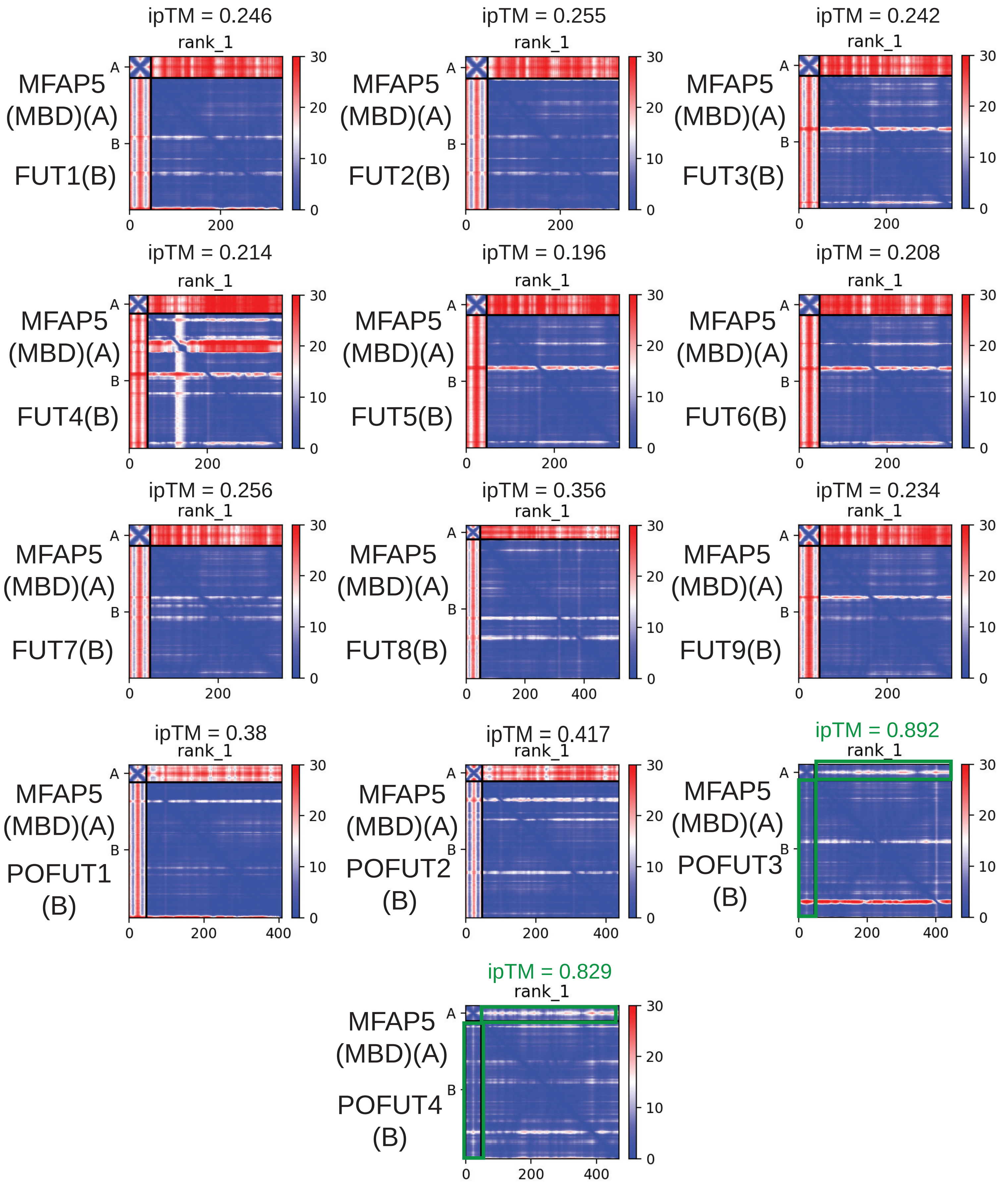


**Supplementary Fig. 5. FUT interaction screen with MFAP5 MBD.** PAE plots for the human MFAP5-fucosyltransferase AlphaFold2-multimer binding assay performed in Colabfold[21]. In each plot the MBD is molecule “A” and the fucosyltransferase is molecule “B”. The top left and bottom right regions of each plot display the predicted aligned error (PAE) within either “A”, or “B”, respectively. The top right and bottom left regions of each plot display the PAE between “A” and “B”. PAE is recorded in Angstroms, and lower values (blue) in the top right and bottom left regions of each plot indicate higher confidence for the interaction. These high confidence interactions are highlighted for POFUT3-MBD and POFUT4-MBD by green boxes. All other fucosyltransferases have very high error in these regions, indicating low confidence for their interaction. ipTM scores are provided for each plot, which is a quantitative measurement of the accuracy of the predicted interaction between the two molecules.

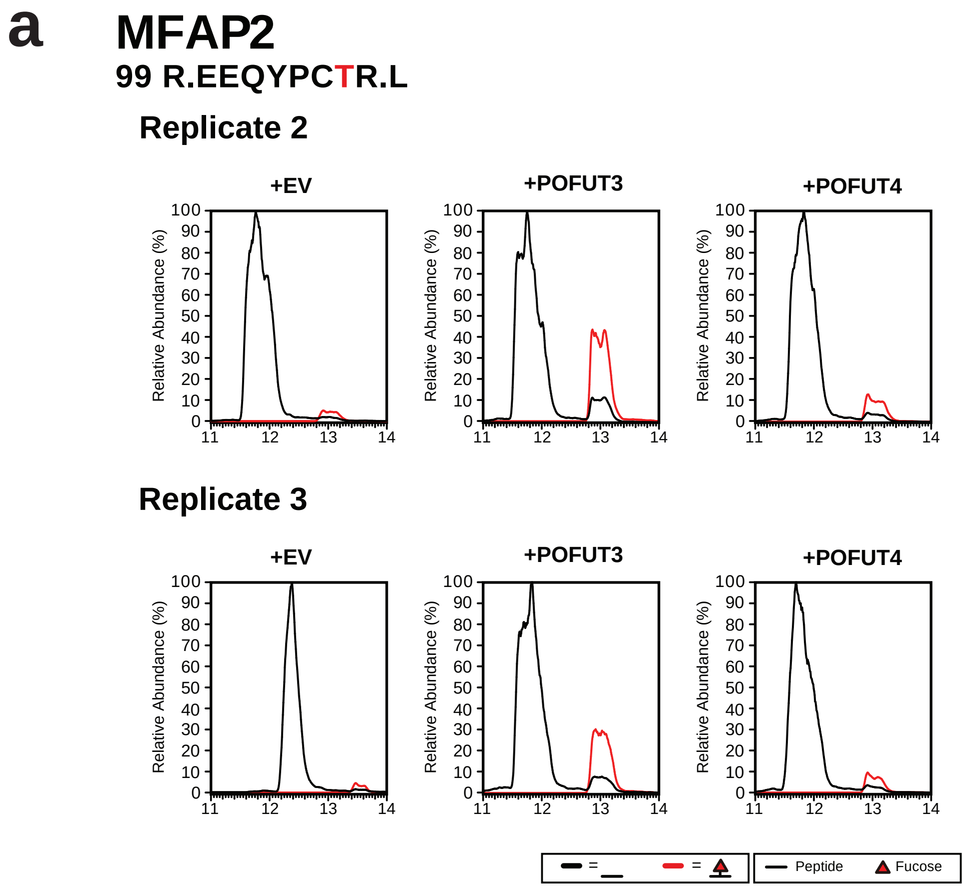


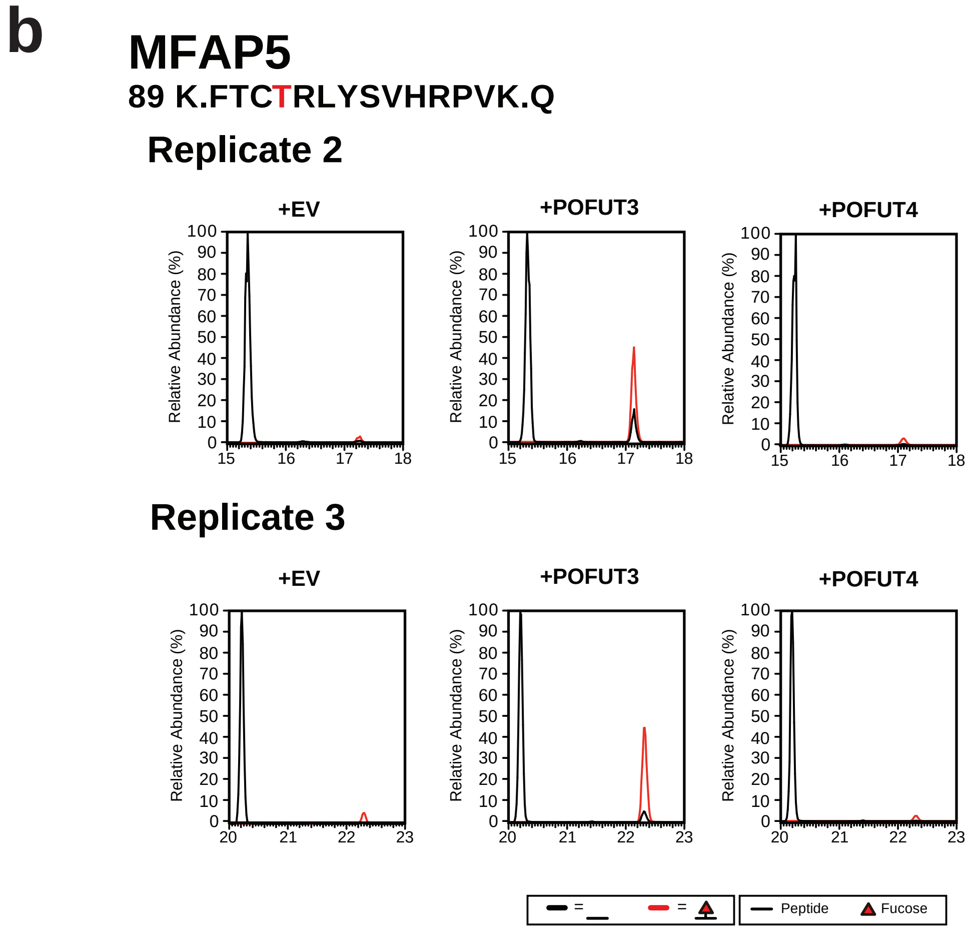


**Supplementary Fig. 6. Additional biological replicates for the data presented in Fig. 2d-g. a**, Two additional biological replicates (n=3, different cultures) for the co-immunoprecipitations of MFAP2 from transfected HEK293 cells. EICs were extracted individually and overlaid to compare the relative intensities of each glycoform. The *O*-fucosylated glycoform is shown in red, while the unmodified glycoform is in black. **b**, Two additional biological replicates (n=3) of the co-immunoprecipitations of MFAP5 from transfected HEK293 cells.

**a**

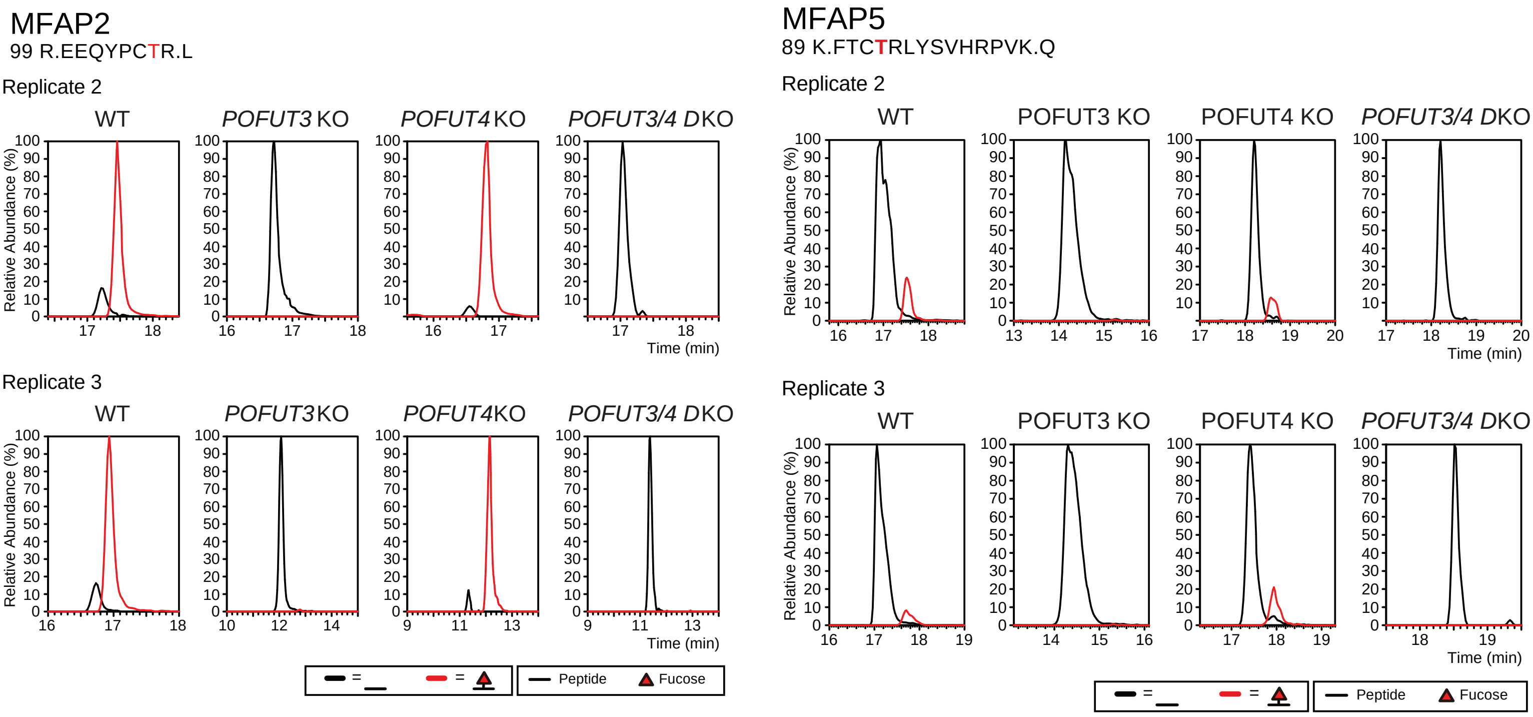


**b**

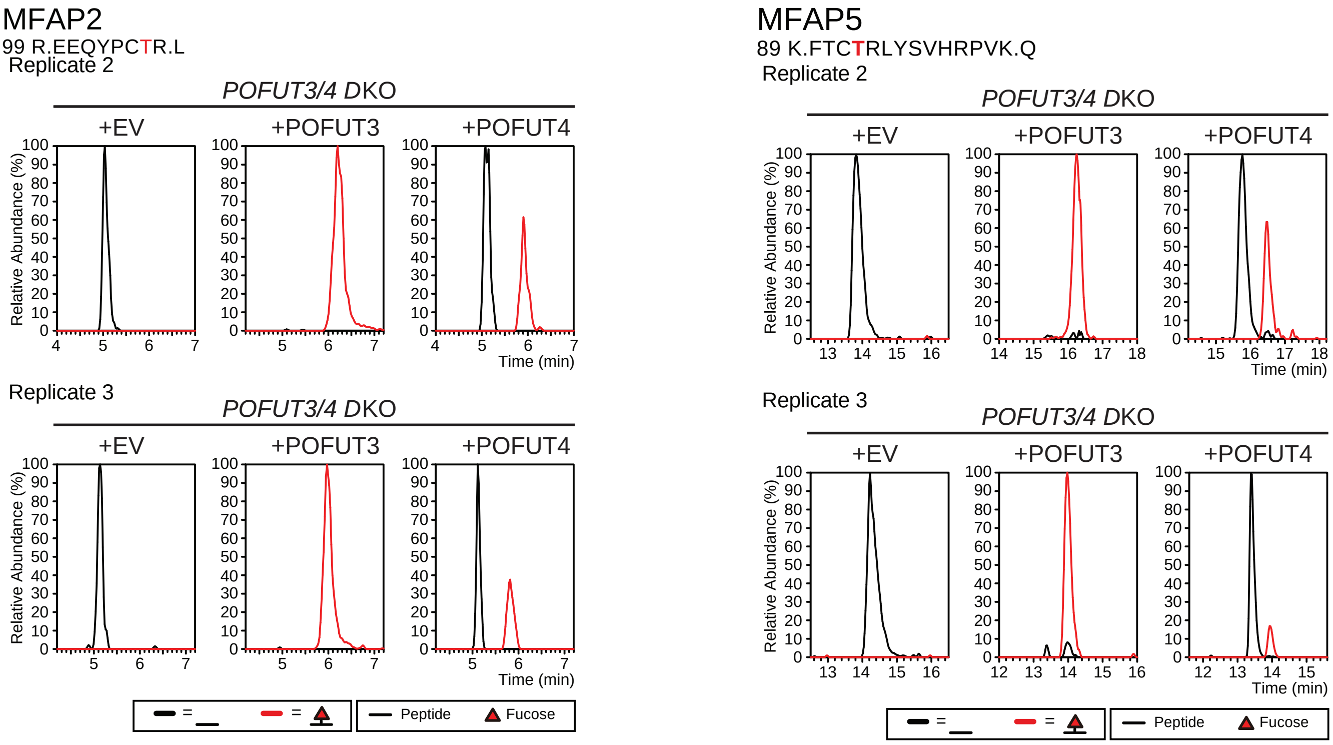


Continued on next page

**c**

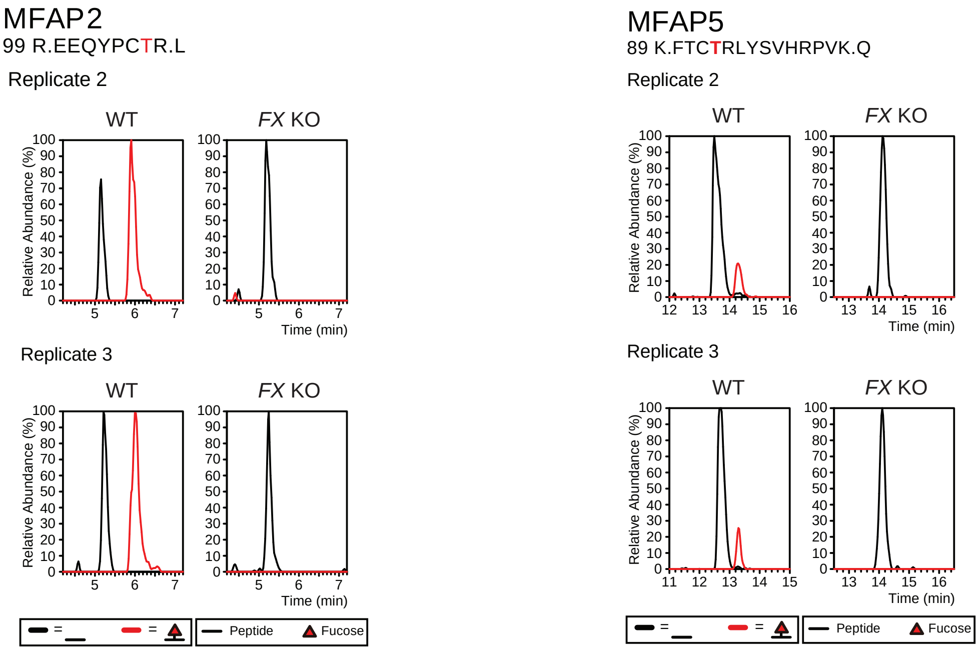


**Supplementary Fig. 7. Additional biological replicates for the data presented in Fig. 3. a**, Two additional biological replicates (n=3, different cultures) for the *POFUT3* KO, *POFUT4* KO and *POFUT3/4* DKO MFAP2/MFAP5 transfections. EICs were extracted individually and overlaid to compare the relative intensities of each glycoform. The *O*-fucosylated glycoform is shown in red, while the unmodified glycoform is in black. **b**, Two additional biological replicates (n=3) of POFUT3/4 rescue assays in *POFUT3/4* DKO cells. **c**, Two additional biological replicates (n=3) of *FX* KO HEK293T transfections comparing *FX* mutant against wildtype.

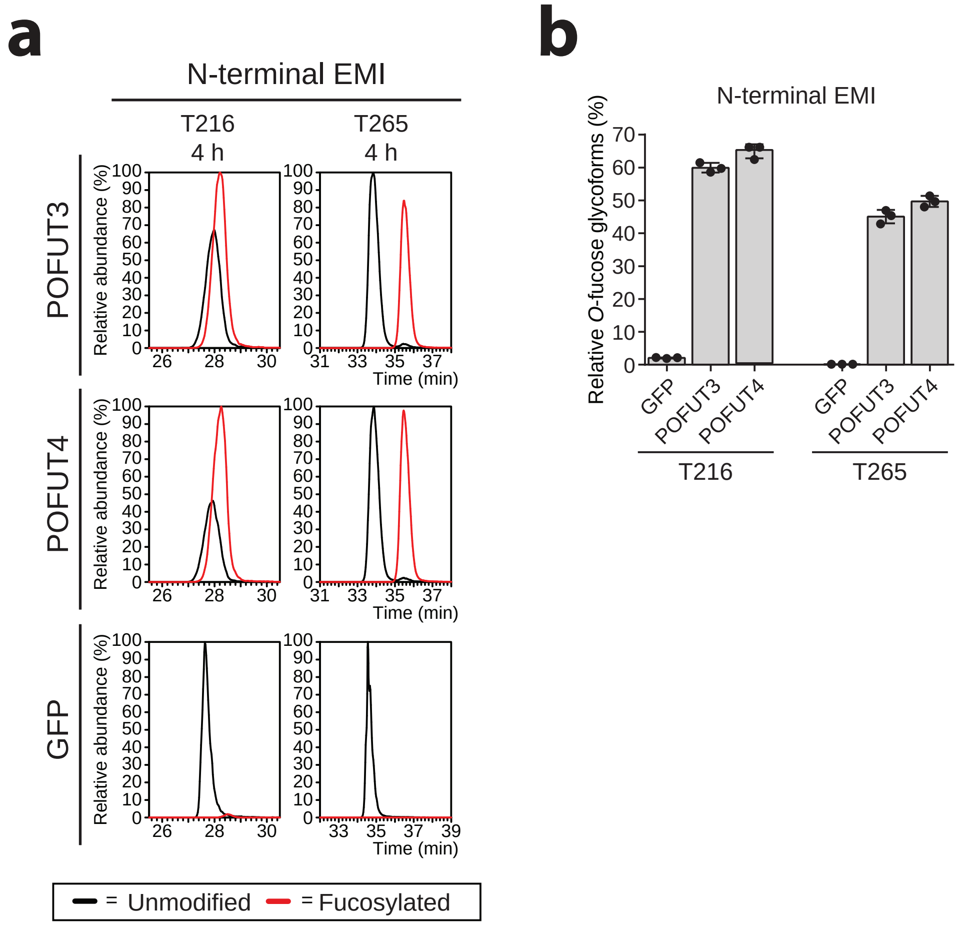


**Supplementary Fig. 8. EMI domain *O*-fucosylation assay controls. a**, *In vitro O*-fucosylation assay where either 0.1 μM purified GFP-POFUT3, GFP-POFUT4, or GFP, was incubated with 0.5 μM non-fucosylated N-terminal EMI domain from MMRN1 and 100 μM GDP-fucose for 4 h. EICs of both the modified and unmodified glycoforms of the peptides containing the T216 and T265 *O*-fucose sites were examined to compare relative abundance after four hours. No significant preference for either POFUT3 or POFUT4 was observed. **b**, Bar graphs show relative abundance of the *O*-fucosylated EMI domain peptides containing the T216 and T265 sites after 4 h incubation.

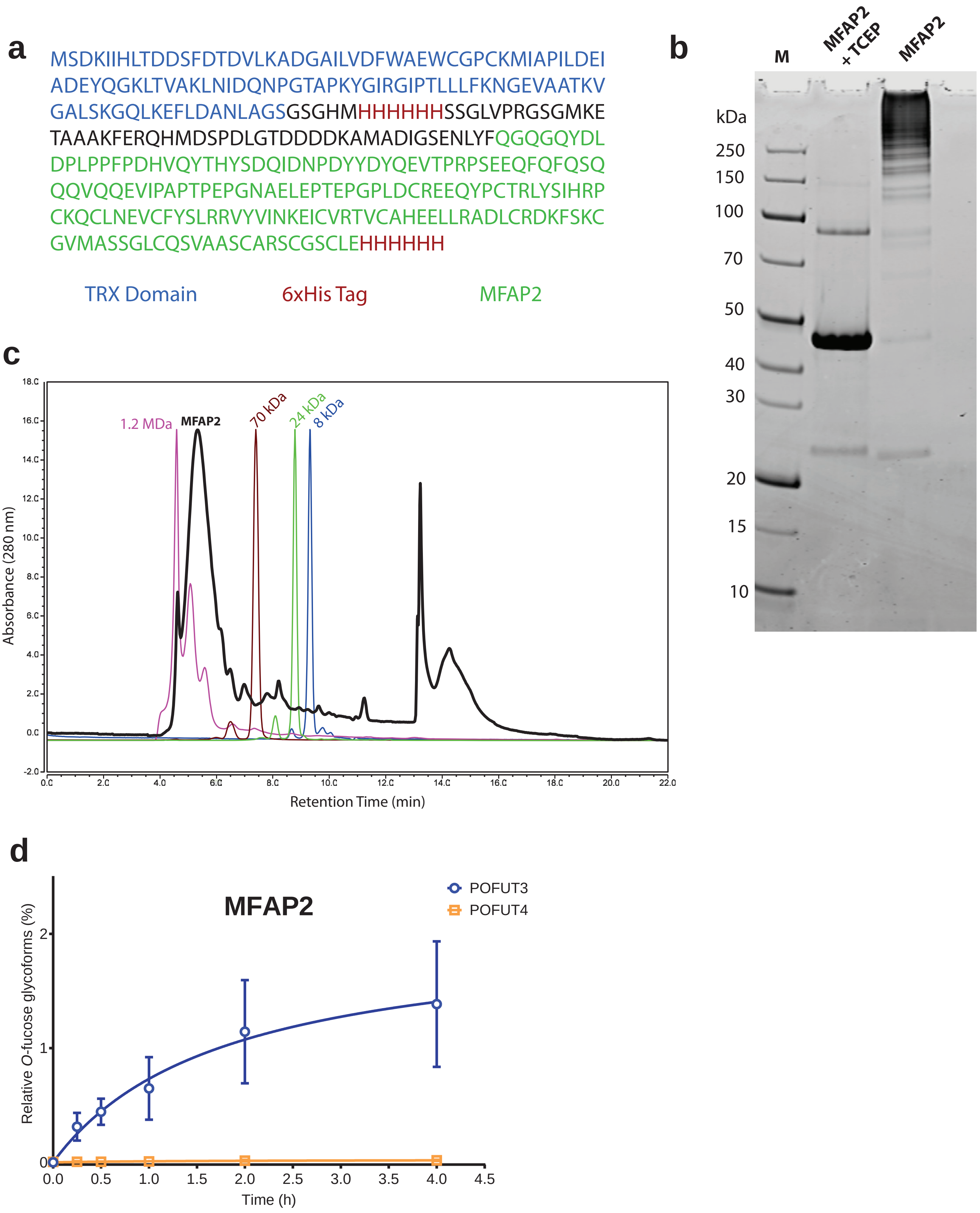


**Supplementary Fig. 9. Bacterial expression of MFAP2 and *in vitro O*-fucosylation assay**. **a**, Protein sequence of the human MFAP2 recombinant protein with TRX and 6xHis tags from the pET-32a vector. **b**, SDS-PAGE and Coomassie stain (total protein) analysis of the purified MFAP2 after bacterial expression, inclusion-body purification and refolding. **c**, Native size exclusion chromatography analysis of the purified MFAP2 protein in PBS buffer compared to several size marker proteins. **d**, *In vitro O*-fucosylation assay where either 0.1 µM purified GFP-POFUT3, or GFP-POFUT4, was incubated with 0.5 µM of bacterially-expressed MFAP2 and 100 µM GDP-fucose for 4 h. Relative abundances of *O*-fucosylation for each reaction were calculated from the EICs of peptides containing the T105 *O*-fucose site and plotted as time-dependent curves. Data are presented as mean ± s.d. from biological triplicates.

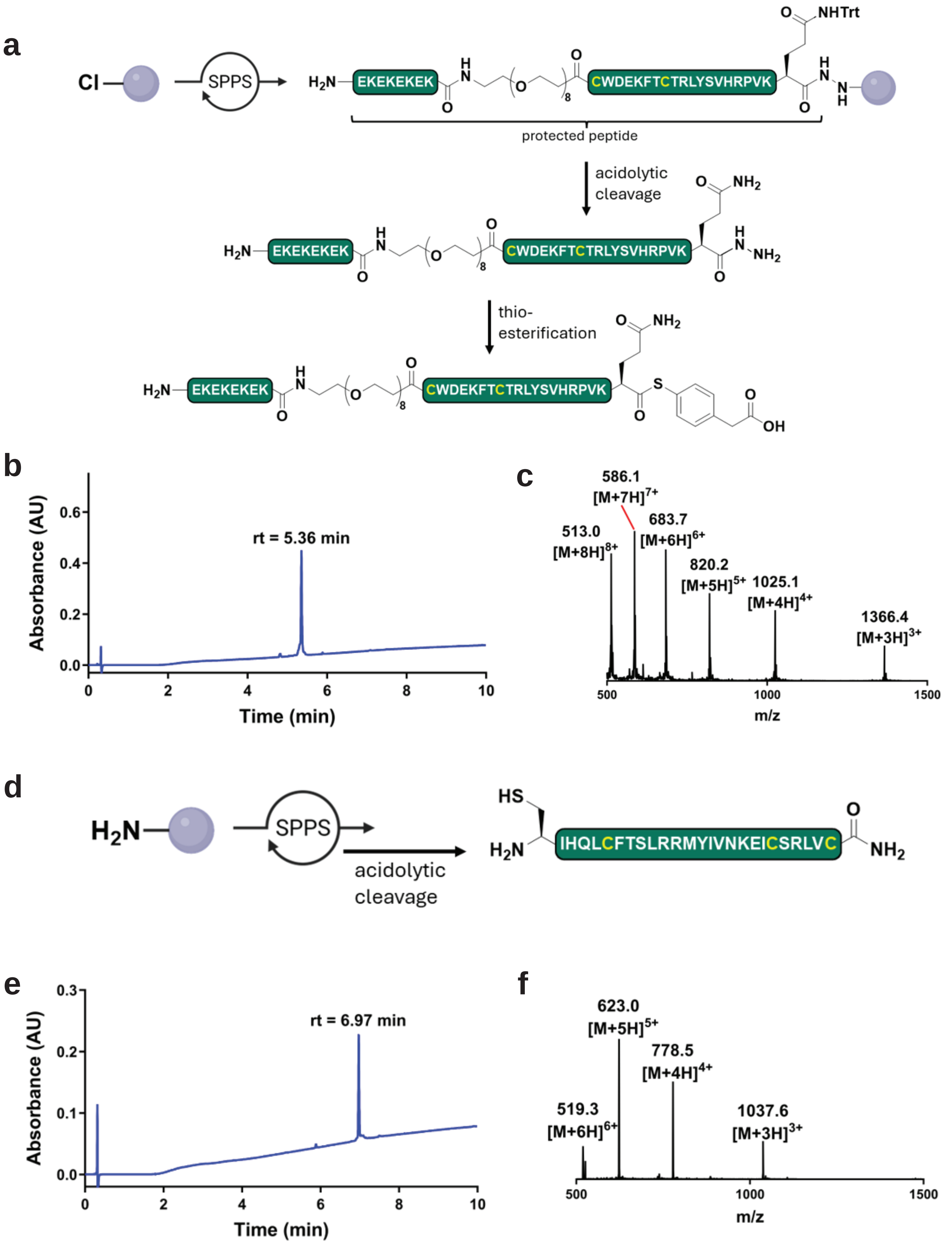


**Supplementary Fig. 10. Synthesis of human MFAP5 matrix binding domain**. **a**, Synthesis of MFAP5-MBD N-terminal thioester fragment bound with solubility tag. **b**, Thioesterification reaction was monitored with UPLC-MS, with a gradient of 0-60% B over 10 min, 0.1% TFA. **c**, ESI-MS of purified peptide thioester upon completion of synthesis. **d**, Synthesis of MFAP5-MBD C-terminal thiol. **e**, MFAP5-MBD C-terminal thiol synthesis UPLC trace with gradient 0-60% B over 10 min, 0.1% TFA. **f**, ESI-MS of purified peptide thiol upon completion of synthesis.

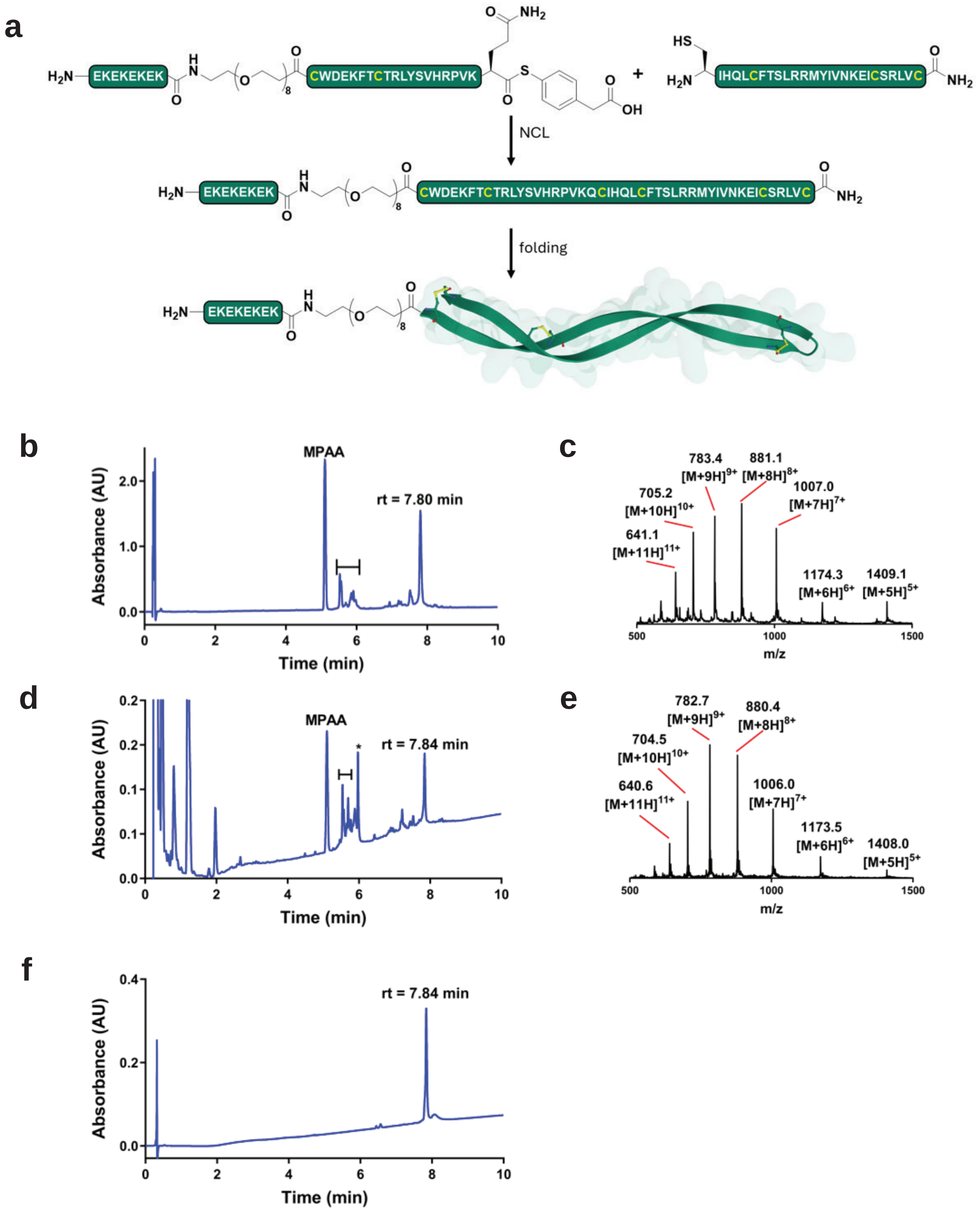


**Supplementary Fig. 11. Native chemical ligation and folding of human MFAP5 matrix binding domain**. **a**, Synthesis and folding of synthetic MFAP5-MBD from ligation of C-terminal and N-terminal peptide fragments. **b**, Crude UPLC trace of ligation reaction with gradient of 0-60% B over 10 min, 0.1% TFA. **c**, ESI-MS of crude reduced ligation product. **d**, UPLC trace of crude folding reaction with gradient of 0-60% B over 10 min, 0.1% TFA. **e**, ESI-MS of purified folded MFAP5. **f**, UPLC trace of purified folded MFAP5 with gradient of 0-60% B over 10 min, 0.1% TFA.

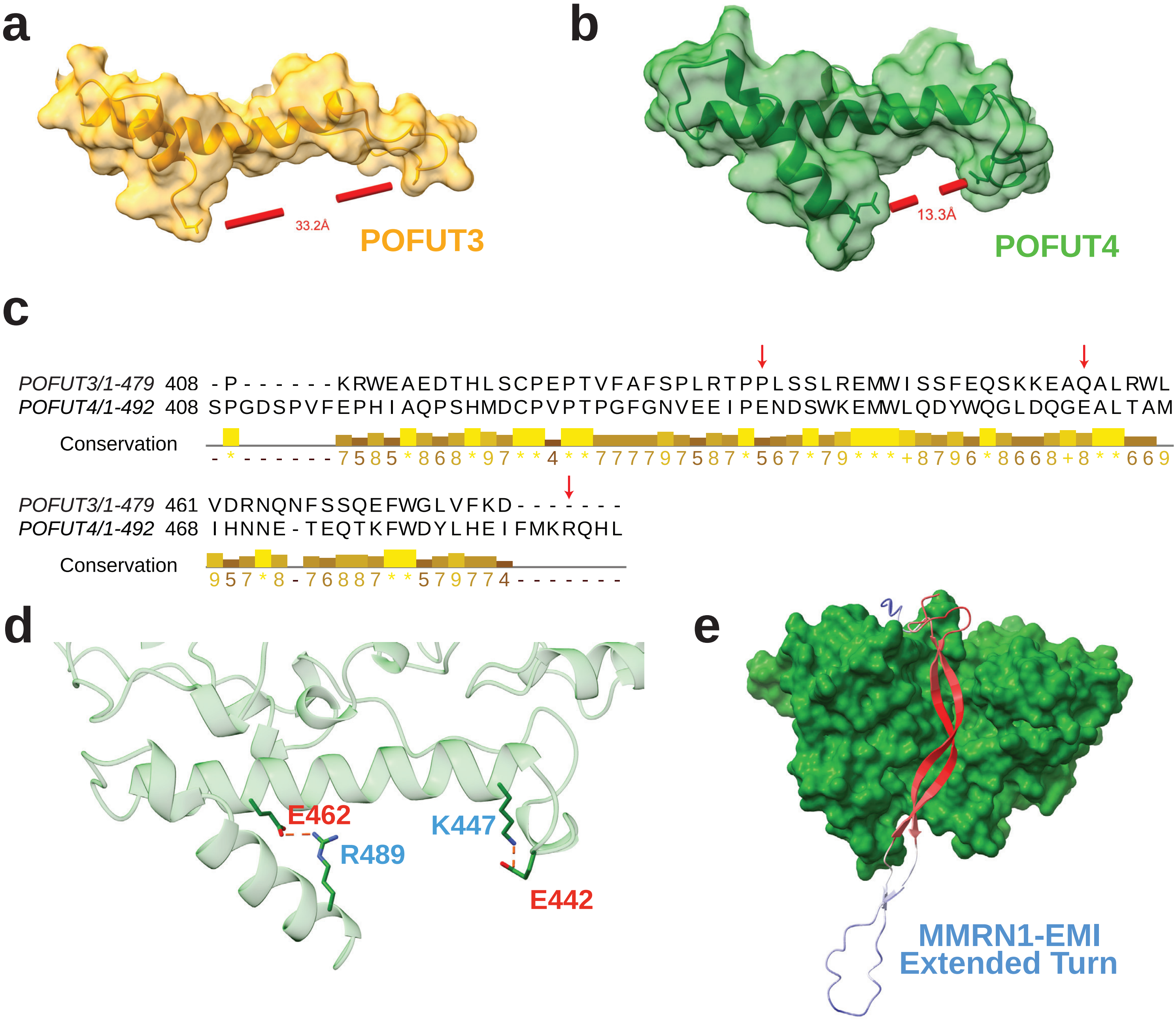


Continued on next page

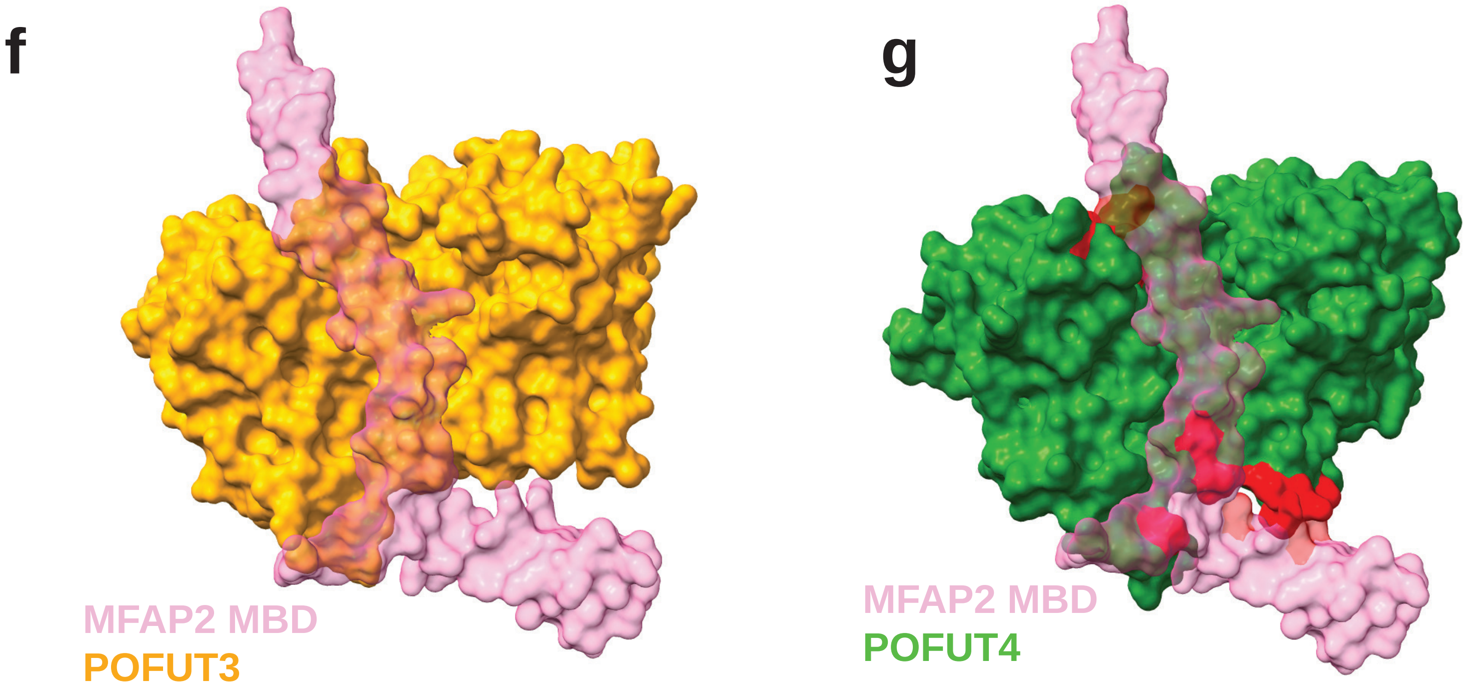


**Supplementary Fig. 12. Analysis of structral clashes in the C-termini of POFUT3/4**. **a**, AlphaFold3 predicted structure for the extended C-terminal region in POFUT3 (residues 415-479), highlighting the open groove structure. **b**, AlphaFold3 predicted structure for the extended C-terminal region in POFUT4 (422-492), highlighting the closed groove structure. **c**, Clustal Omega alignment of human POFUT3 and POFUT4 C-terminal regions. Arrows highlight putative salt bridge-forming residues unique to POFUT4. d, Predicted salt bridge-forming residues in the predicted POFUT4 C-terminus structure. **e**, MMRN1's EMI domain features an extended beta turn structure, likely imparting flexibility to bind either POFUT3 or POFUT4. **f-g**, Alphafold2-multimer predicted structure for the interaction of MFAP2's MBD with POFUT3 (**f**) or POFUT4 (**g**) with predicted steric clashes highlighted in red. Clashes predicted by ChimeraX (v. 1.11.1) "Clashes" algorithm using default settings.

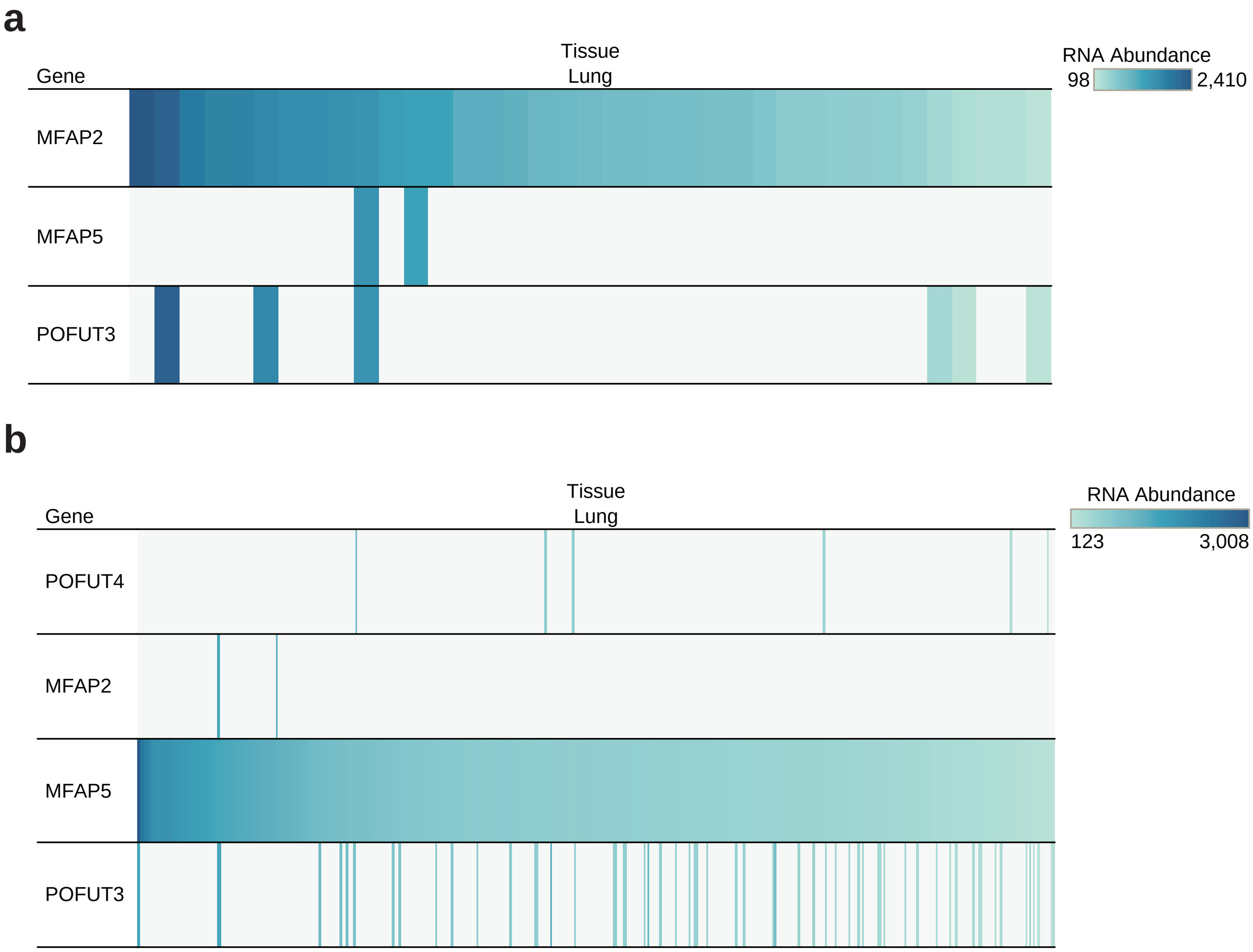


**Supplementary Fig. 13. GTEx Single cell RNA dataset for human lung tissue**. **a**, Human lung single-cell RNA sequencing data from GTEx[31] indicating the patterns of MFAP2 mRNA expression in comparison to MFAP5, POFUT3 and POFUT4 (not detected). **b**, Human lung single-cell RNA sequencing data from GTEx indicating the patterns of MFAP5 mRNA expression in comparison to MFAP2, POFUT3 and POFUT4.

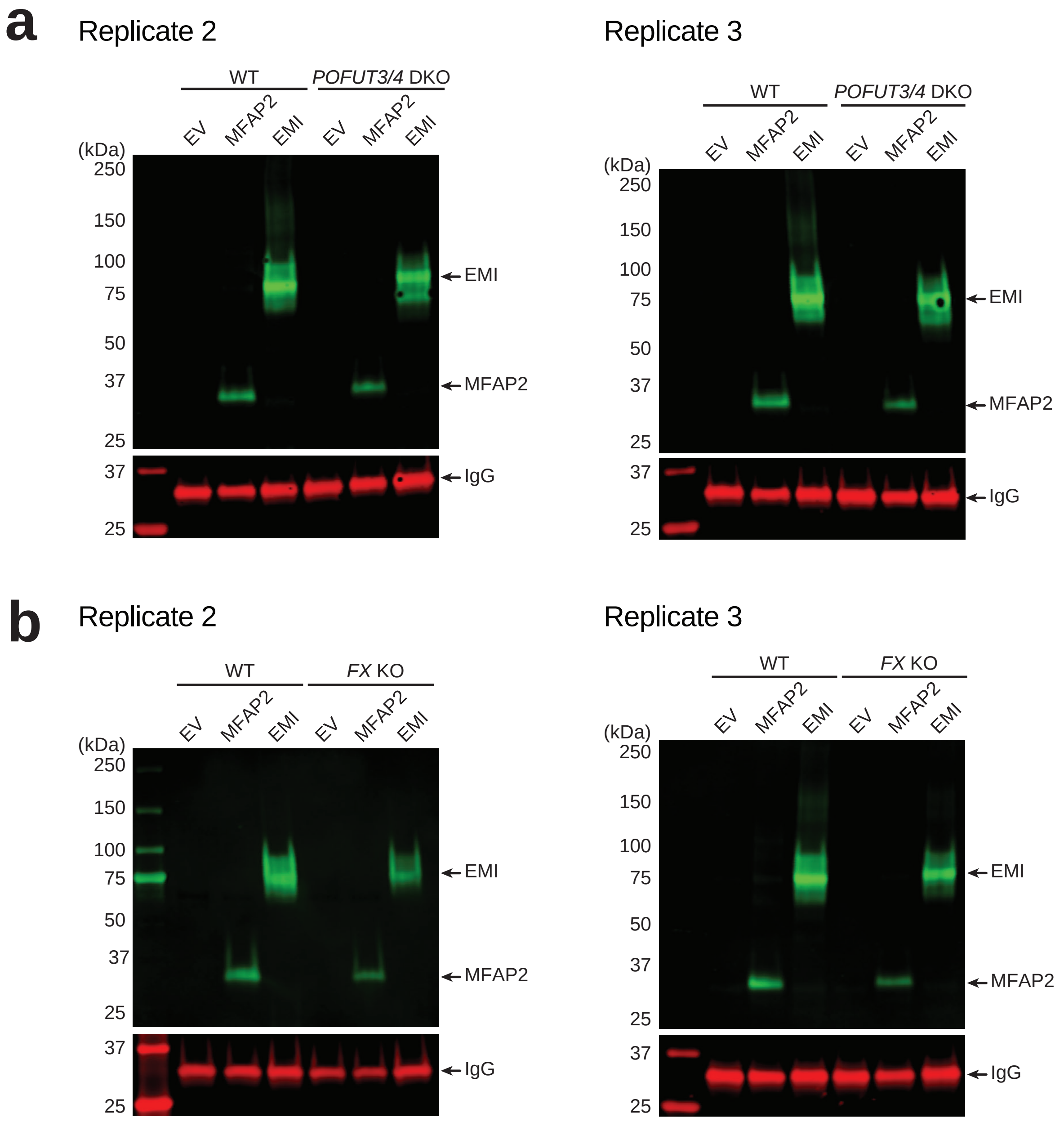


**Supplementary Fig. 14. Additional biological replicates for the data presented in Fig. 5e,f. a**, HEK293T WT or *POFUT3/4* KO cells were transfected with plasmids encoding Myc-tagged MFAP2, N-terminal EMI (positive control) or EV, alongside an IgG secretion control. Two-day culture media was analysed by western blot probed with anti-Myc and anti-human IgG antibodies. **b**, HEK293T WT or *FX* KO cells were transfected with plasmids encoding Myc-tagged MFAP2, N-terminal EMI or EV, alongside an IgG secretion control. Two-day culture media was analysed by western blot probed with anti-Myc and anti-human IgG antibodies.

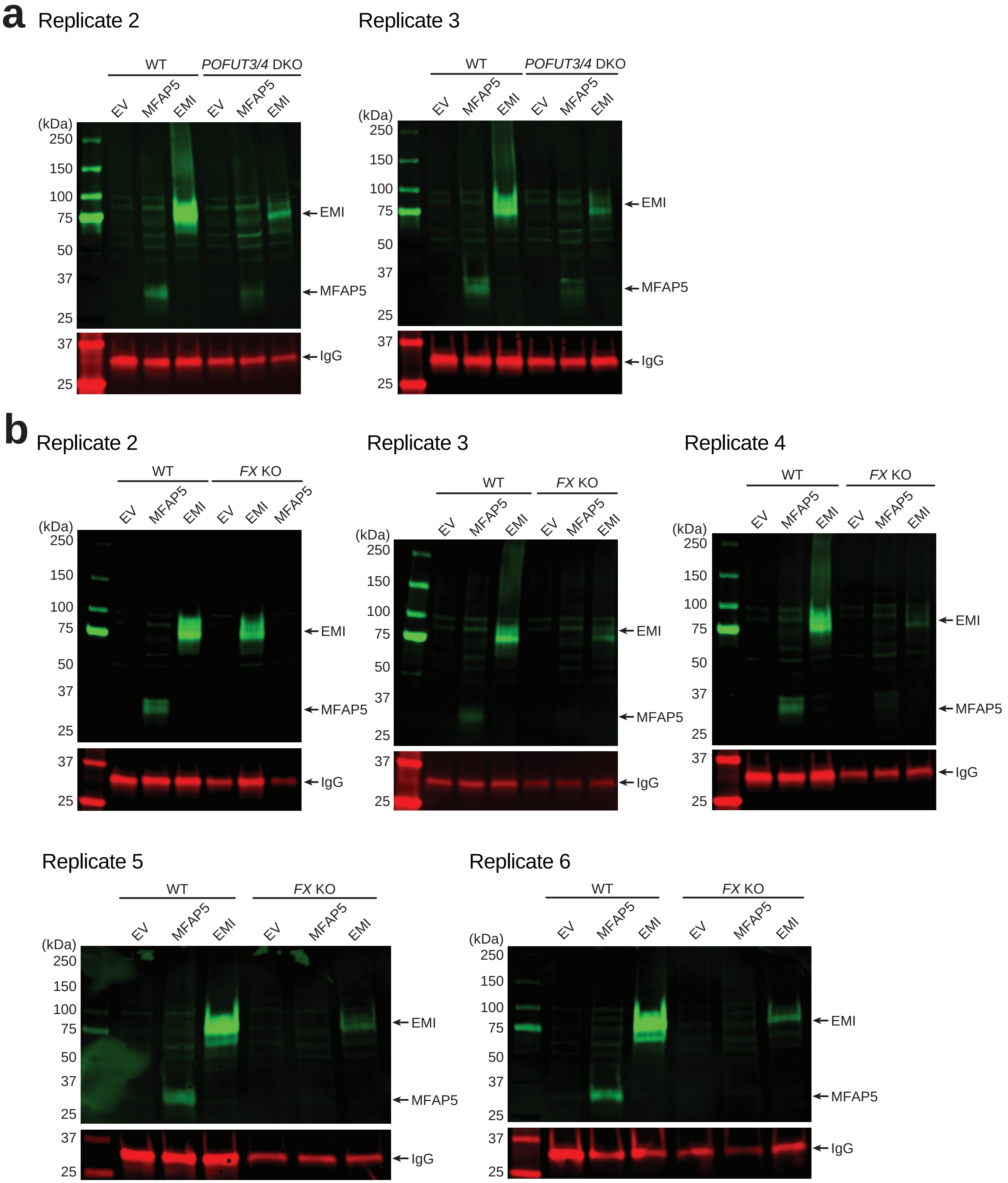


**Supplementary Fig. 15. Additional biological replicates for the data presented in Fig. 5i,j. a**, HEK293T WT or *POFUT3/4* KO cells were transfected with plasmids encoding Myc-tagged MFAP5, N-terminal EMI (positive control) or EV, alongside an IgG secretion control. Two-day culture media was analysed by western blot probed with anti-Myc and anti-human IgG antibodies. **b**, HEK293T WT or *FX* KO cells were transfected with plasmids encoding Myc-tagged MFAP5, N-terminal EMI or EV, alongside an IgG secretion control. Two-day culture media was analysed by western blot probed with anti-Myc and anti-human IgG antibodies.
