## Supplementary Methods 1 for "Iterative structural homology search identifies new substrates of the protein *O*-fucosyltransferases POFUT3 and POFUT4"

### Supplementary Methods – MFAP5 Protein Synthesis and Folding

**Materials:** Reagents were used as received unless otherwise noted. Resins, coupling reagents, and amino acids were all obtained from GL Biochem, Mimotopes, or Novabiochem. *N*,*N*-dimethylformamide (DMF) was obtained as peptide synthesis grade from Merck or Labscan. All Gdn·HCl used to prepare buffers was dried *in vacuo* before use.

**C8 UPLC Column:** Waters Acquity BEH, 130Å, 1.7 µm, 2.1 mm × 50 mm.

UPLC was performed with a flow rate of 0.2 mL/min and a mobile phase composed of water (solvent A) and acetonitrile (solvent B) with 0.1 vol.% TFA as the additive.

The analysis of all chromatograms was conducted using Empower 3 Pro software (2010).

**Preparative HPLC:** Preparative reverse-phase HPLC was performed using a Waters 600 Multisolvent Delivery System and Waters 500 pump with 2996 photodiode array detector or Waters 490E Programmable Wavelength Detector operating at 214 and 280 nm. Peptide fragments and ligated products were purified using the following columns as described in the experimental sections:

**C8 Semi-Preparative HPLC Column:** XBridge BEH, 300 Å, 5 µm, 40 mm × 160 mm.

**UPLC-MS:** UPLC-MS was performed on a Shimadzu LC-MS 2020 system equipped with a Nexera X2 LC-30AD pump and a Nexera X2 SPD-M30A diode array detector coupled to a Shimadzu 2020 electrospray ionization-mass spectrometer (ESI-MS) operating in positive mode. Peptides and proteins were analyzed using a C8 UPLC Column (see above) with a gradient of 0-100% B over 3 min, 0.1% TFA. All mass spectra extracted from UPLC-MS data are from the total ion count of the entire UPLC gradient.

**Ligation and folding yields:** All ligation and folding yields were calculated using an analytical balance and verified through nanodrop. Yields were **not** compensated for aliquots removed for UPLC-MS analysis.

**Loading 2-chlorotrityl chloride resin:** 2-chlorotrityl chloride resin (1.22 mmol/g loading) was swollen in $CH_2Cl_2$ for 30 min. A solution of Fmoc-hydrazine or Fmoc-protected amino acid (2 eq. relative to maximum loading) and *N,N*-di*iso*propylethylamine (DIPEA, 8 eq.) in $CH_2Cl_2$ (0.3 M of Fmoc-hydrazine) was added to the resin and agitated for 16 h at rt. The loading solution was drained and the resin was rinsed with DMF (5 × 3 mL), $CH_2Cl_2$ (5 × 3 mL), and DMF (5 × 3 mL). The resin was treated with a capping solution of $CH_2Cl_2$/MeOH/DIPEA (17:2:1 v/v/v, 4 mL) for 1 h then washed with DMF (5 × 3 mL), $CH_2Cl_2$ (5 × 3 mL), and DMF (5 × 3 mL).

**Quantification of resin loading:** 10 mg of the loaded resin was treated with a solution of 2 vol.% 1,8-diazabicyclo[5.4.0]undec-7-ene (DBU) in DMF (2 mL) and agitated for 30 min. The solution was removed from resin and diluted to 10 mL with acetonitrile. A 1 mL aliquot of this resultant solution was taken up and diluted to 12.5 mL with acetonitrile. The absorbance of the DBU-fulvene adduct ($\lambda$ = 304 nm, $\varepsilon$ = 9254 $M^{-1}$ $cm^{-1}$) was measured to estimate the resin loading.

**General method using SYRO I Automated Synthesizer:** Heated automated Fmoc-SPPS was performed on a Biotage SYRO I automated synthesizer on a 50 µmol scale. Solutions of Fmoc-AA-OH (0.5 M) and Oxyma (0.55 M) in DMF, and *N,N'*-di*iso*propylcarbodiimide (DIC, 0.5 M) in DMF were prepared for coupling. General synthetic protocols for each step in the Fmoc-SPPS cycle are provided below:

*Fmoc deprotection*: The resin-bound peptide was shaken in a solution of 40 vol. % piperidine in DMF (800 µL, 4 min). The deprotection solution was then drained and the resin was treated again with a solution of 20 vol. % piperidine in DMF (800 µL, 4 min). The deprotection solution was then drained and the resin was washed with DMF (4 × 1.0 mL).

*Coupling*: The resin-bound peptide was shaken in a solution of Fmoc-AA-OH (4 eq., 0.17 M), Oxyma (4.4 eq., 0.18 M) and DIC (4 eq., 0.17 M) in DMF for 45 min at 40 °C. The coupling solution was drained, and the resin was retreated with a fresh coupling solution then washed with DMF (4 × 800 µL).

*Capping*: The resin-bound peptide was shaken in a solution of 5 vol. % $Ac_2O$ and 10 vol. % DIPEA in DMF (800 µL, 6 min). The capping solution was then drained, and the resin was washed with DMF (4 × 800 µL).

**Acidolytic cleavage from resin with concomitant side-chain deprotection:** Resin was initially washed with $CH_2Cl_2$ (2 × 3 mL), then a mixture of trifluoroacetic acid (TFA), tri*iso*propylsilane (TIS) and water (18:1:1 v/v/v, 5 mL) was added to the resin-bound peptide (50 µmol) and agitated at rt for 2 h. The resin was then filtered and washed with TFA (2 × 3 mL) and the combined filtrates were concentrated under nitrogen flow. Diethyl ether (40 mL) was added and the suspension cooled to 0 °C for 10 min. The precipitate was pelleted by centrifugation at 6800 rcf for 10 min at 0 ˚C and the supernatant decanted.

**MFAP5 N-terminal thioester fragment bound with solubility tag**

Fmoc-hydrazine was loaded onto 2-CTC resin in accordance with the general methods. The loaded resin (25 μmol) was subjected to Fmoc-SPPS as outlined in the general methods (see SYRO I automated synthesizer section) where the final amino acid was Fmoc deprotected and the resin was washed as outlined in the general methods (see SYRO I automated synthesizer section). Acidolytic cleavage as described in general methods was then conducted and the crude peptide acyl hydrazide was lyophilized to help with solubility for the next step (**Supplementary Figure 11a**). A buffer solution containing 100 mM TCEP, 200 mM HEPES, 50 mM MPAA and 6 M Gnd.HCl, pH 2.5 was used to dissolve the peptide acyl hydrazide to a final concentration of 5 mM, followed by the addition of 15 eq. of acetylacetone (acac). The solution was then allowed to stir at room temperature for 2-3 h. Completion of the thioesterification reaction was monitored by UPLC-MS (**Supplementary Figure 11b, c**). Upon completion the crude reaction mixture was subjected to HPLC purification and lyophilization to afford the purified peptide thioester as a white fluffy solid (6 mg, 6% yield).

**MFAP5 C-terminal thiol**

Rink-amide resin (25 μmol) was subjected to Fmoc-SPPS as outlined in the general methods (see SYRO I automated synthesizer section) where the final amino acid was Fmoc deprotected and the resin was washed as outlined in the general methods (see SYRO I automated synthesizer section). Acidolytic cleavage as described in general methods was then conducted (**Supplementary Figure 11d**) and the crude peptide was subjected to HPLC purification and lyophilization to afford the purified peptide thiol as a white fluffy solid (11 mg, 14% yield) (**Supplementary Figure 11e, f**)

**Folded MFAP5 with solubility tag**

Two separate solutions of MFAP5 thioester fragment (3.0 mg, 12 mM, 1.2 equiv.) and MFAP5 thiol fragment (1.8 mg, 10 mM, 1 equiv.) were made by dissolving the respective lyophilized peptide fragments in ligation buffer comprising 6 M Gdn·HCl, 1 M HEPES, 200 mM TCEP, and 50 mM MPAA at pH 6.5 (50 µL each) and immediately mixed and shaken on an orbital shaker for 2 hours at 37 ºC (**Supplementary Figure 12a**). After complete ligation was observed (by UPLCMS) (**Supplementary Figure 12b, c**), the crude ligation solution was subjected to a stepwise dialysis folding procedure. Specifically, the ligation mixture diluted 1 in 10 with 6 M Gdn·HCl, 0.1 M HEPES, 10 mM DTT at pH 6.8 (900 µL) and left to stand for 10 min to produce the completely reduced MFAP5 ligation product. The solution was then diluted equivolume with buffer comprising of 2 M Gdn·HCl, 50 mM Tris, 0.5 M NaCl and 1 M L-Arginine at pH 7.5. The sample was then added to snakeskin dialysis tubing (3.5 kDa cutoff) and placed in 1 L of buffer comprising 2 M Gdn·HCl, 50 mM tris, 0.5 M NaCl and 0.5 M L-Arginine, 3 mM GSH and 1 mM GSSH at pH 7.5 for 16 h at 4 ºC. This led to conversion to a folded MFAP5 with 3 disulfide bridge as determined by UPLCMS and a shift in retention time in analytical UPLC (**Supplementary Figure 12 d-f**). Following purification by HPLC, folded MFAP5 was isolated as a fluffy white solid (0.5 mg, 12% yield).
